# HemK2 functions for sufficient protein synthesis and RNA stability through eRF1 methylation during *Drosophila* oogenesis

**DOI:** 10.1101/2024.01.29.576963

**Authors:** Fengmei Xu, Ritsuko Suyama, Toshifumi Inada, Shinichi Kawaguchi, Toshie Kai

## Abstract

HemK2 is a highly conserved methyltransferase spanning from yeast to humans. Despite its conservation, the identification of its genuine substrates has been controversial, and its biological importance in higher organisms remains unclear. In this study, we elucidate the role of HemK2 in the methylation of eukaryotic Release Factor 1 (eRF1), a process essential for female germline development in *Drosophila melanogaster*. Knockdown of *hemK2* in the germline cells (*hemK2*-GLKD) induces apoptosis in these cells, accompanied by a pronounced decrease in both eRF1 methylation and protein synthesis. The overexpression of a methylation-deficient eRF1 variant recapitulates the defects observed in *hemK2*-GLKD, suggesting that eRF1 is a primary methylation target of HemK2. Furthermore, *hemK2*-GLKD leads to significant reduction mRNA levels in germline cell. We demonstrate that these defects in oogenesis and protein synthesis can be partially restored by inhibiting the No-Go Decay pathway. In addition, *hemK2* knockdown is associated with increased disome formation, suggesting that disruptions in eRF1 methylation may provoke ribosomal stalling, which subsequently activates translation-coupled mRNA surveillance mechanisms that degrade actively-translated mRNAs. We propose that HemK2-mediated methylation of eRF1 is critical for ensuring efficient protein production and mRNA stability, which are vital for the generation of high-quality eggs.

## Introduction

Post-translational modifications (PTMs), such as methylation, phosphorylation, glycosylation, acetylation, and ubiquitination, are chemical alterations that occur after protein synthesis. PTMs profoundly influence protein function, interaction, stability, and enzymatic activity (Ramazi and Zahiri, 2021; Ryšlavá et al., 2013). Among the over 400 types of PTMs, methylation is one of the most pervasive. Methylation reactions are catalyzed by S-adenosylmethionine-dependent methyltransferases (MTases), which transfer methyl groups to a variety of biomolecules, including proteins, DNA, and RNA, targeting cytosines and adenines (Chen et al., 2013; Dai et al., 2021; Ping et al., 2014; Wang et al., 2016). In the *Drosophila* genome, 127 MTase genes have been annotated, and these MTases have been further classified based on their substrate specificities. Lysine MTases and arginine MTases constitute substantial groups among protein MTases. Prior studies on histone methylation have highlighted the importance of lysine/arginine modifications in transcriptional regulation (Gates et al., 2017; Zhao and Garcia, 2015). In addition to lysine and arginine residues, other amino acid residues such as histidine (Clarke, 2013), aspartic acid (Sprung et al., 2008), asparagine (Klotz et al., 1990), and glutamine (Heurgué-Hamard et al., 2002) are also subject to methylation.

HemK2 is a member of the unclassified protein MTases, distinct due to its broad substrate class. HemK2 has been previously shown to catalyze the methylation of histone H4 lysine 12 (H4K12), DNA N6-adenine (6mA), and eukaryotic translation release factor 1 (eRF1). HemK2-mediated H4K12 methylation is implicated in regulating the proliferation of prostate tumor cells (Metzger et al., 2019), while diminished genomic DNA 6mA levels have been associated with tumorigenesis in cancer patients (Xiao et al., 2018). However, the enzymatic activity of HemK2 towards genomic DNA 6mA remains controversial (Kweon et al., 2019; Ratel et al., 2006; Woodcock et al., 2019). Conversely, HemK2’s role as an MTase for eRF1 has been extensively investigated in yeast and mammalian cells. HemK2 (N6AMT1 in humans; YDR140w or Mtq2p in yeast) methylates the glutamine residue within the GGQ motif of eRF1 (Figaro et al., 2008; Heurgué-Hamard et al., 2006; Liu et al., 2010). Depletion of HemK2 in yeast (Mtq2p) resulted in sensitivity to translation-fidelity antibiotics (Polevoda et al., 2006). In mice, HemK2 (also known as N6AMT1) deficiency led to early embryonic lethality and impaired post-implantation development (Liu et al., 2010). Yet, the methylation of these potential substrates by HemK2 under physiological conditions in multicellular organisms is not well defined.

*Drosophila* oogenesis serves as an exemplary model for studying the genetic regulation of egg production. The female fly harbors a pair of ovaries, each comprising 18–20 ovarioles, which contain the germarium at the anterior tip, succeeded by sequentially maturing egg chambers (Avilés-Pagán and Orr-Weaver, 2018; Gleason et al., 2018; Rastegari et al., 2020). Each egg chamber encompasses one oocyte and 15 nurse cells that support oocyte growth, surrounded by a monolayer of somatic follicle cells. In nurse cells, the production of mRNAs and proteins for maternal deposition into eggs is critical for proper oogenesis. Consequently, *Drosophila* oogenesis is an energetically demanding process involving cellular proliferation and growth that necessitates extensive protein synthesis (Lasko, 2012).

The role of HemK2 as a methyltransferase (MTase) has been extensively investigated in yeast and mammalian cell cultures. Nonetheless, its function within multicellular organisms, particularly in the reproductive system—characterized by active gene expression and protein synthesis—has not been fully elucidated. In this study, we delineate the role of HemK2 in eRF1 methylation during *Drosophila* oogenesis. Our findings indicate that HemK2 is critical for egg production; knockdown of *hemK2* expression arrested oogenesis at an intermediate stage and induced apoptosis in germline cells. The disruption of *hemK2* expression resulted in a substantial decrease in eRF1 methylation, whereas methylation of DNA 6mA and histone H4 lysine 12 (H4K12) was not significantly affected. Moreover, the lack of eRF1 methylation impaired translation efficiency, which in turn reduced mRNA levels via the No-Go Decay pathway. Collectively, these results highlight the indispensable function of HemK2-mediated eRF1 methylation in promoting effective protein synthesis and mRNA stability, thus modulating gene expression throughout *Drosophila* oogenesis.

## Results

### Germline knockdown of *hemK2* resulted in female sterility marked by the developmental arrest in the mid-oogenesis

To investigate the role of HemK2, a highly conserved methyltransferase encoded by *CG9960* in *Drosophila melanogaster*, we employed RNA interference to repress its expression. This knockdown was conducted by shRNA targeted to the 3’UTR (BL#40837) using the germline-specific driver *NGT40*; *NosGal4-VP16* (Grieder et al., 2000). In normal development, the oocyte accrues proteins and RNAs from 15 nurse cells through cytoplasmic channels (Fig. 1A, B). Females with germline-specific knockdown of *hemK2* (*hemK2*-GLKD) showed a ∼65% decrease in mRNA levels (Fig. 1C, S1A) and exhibited sterility. These *hemK2*-GLKD females contained small ovaries and were incapable of laying eggs (Fig. 1B, D), indicative of developmental arrest at stages 7/8 before the vitellogenic stages. Conversely, *hemK2*-GLKD males, despite a similar reduction in its mRNA level to approximately 25%, did not exhibit notable fertility impairments (Fig. S1F, G).

**Figure 1:**
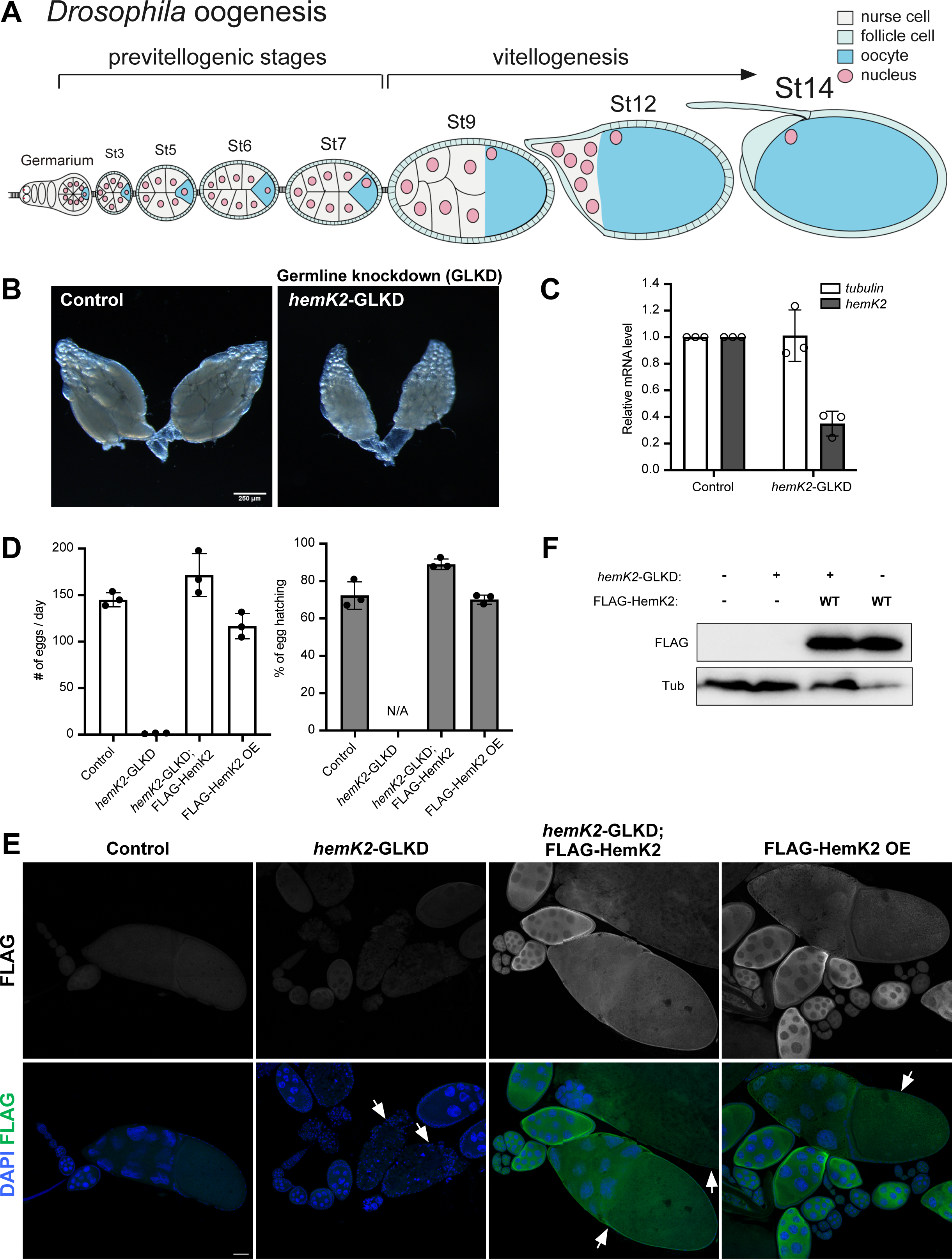
HemK2 is essential for *Drosophila* germline development. (A) Diagrammatic illustration of *Drosophila* oogenesis. Each ovariole is composed of a germarium at the anterior end, followed by progressively developing egg chambers. A 16-cell germline cyst enclosed with somatic follicle cells buds out of germarium, forming an egg chamber. Of those 16 germline cells, one differentiates into the oocyte, while others become nurse cells. (B) Representative images of ovarian morphology from control and *hemK2* germline knockdown (GLKD) samples. *hemK2*-GLKD leads to female sterility, characterized by arrested development of egg chambers at stages 7-8 and impaired vitellogenesis. Scale bar, 250 µm. (C) Quantitative RT-PCR confirmation of knockdown efficacy in *hemK2*-GLKD ovaries. Expression levels are normalized to *rp49* with *tubulin* serving as an internal control. Standard deviation is denoted by error bars (n=3). (D) Analysis of egg laying and hatching rate. Infertility observed in *hemK2*-GLKD is rescued by expression of the wild-type FLAG-tagged HemK2 transgene using the germline-specific driver *NGT40*; *nosGal4-VP16*. Daily egg production by 3 females was recorded (n=3). Error bars represent standard deviation. (E) Immunofluorescence of FLAG-tagged HemK2 (green) and DNA (DAPI, blue) in ovaries of the indicated genotypes corresponding to (D). Arrows highlight degenerated egg chambers in *hemK2*-GLKD and later-stage egg chambers following FLAG-HemK2 expression. Scale bars, 50 µm. (F) Western blot analysis detecting FLAG epitope to demonstrate germline expression of FLAG-HemK2 wildtype in ovarian extracts. Tubulin is utilized as a loading control.

HemK2 is expressed from a dicistronic transcript which is co-transcribed with the upstream open reading frame, *snapin*, in a single mRNA (Fig. S1A) (Wall et al., 2005). To confirm that the observed defects in *hemK2*-GLKD were due to diminished HemK2 protein expression, individual transgenes coding for N-terminal FLAG-tagged HemK2 or Snapin, lacking the shRNA target sequence, were expressed in the germline following knockdown of the dicistronic mRNA (Fig. 1E; Fig. S1E). Females expressing the *hemK2* transgene alone in their germline exhibited egg laying and hatching rates comparable to controls (Fig. 1D), thus rescuing the sterility defect. In contrast, expression of *snapin* did not restore these defects (Fig. S1D). These results confirm that the sterility observed was specifically due to compromised *hemK2* function resulting from knockdown of the dicistronic transcript, establishing *hemK2*’s vital role in germline cells during oogenesis.

To investigate the function of *hemK2* in somatic cells, we knocked down its expression in ovarian somatic cells, including follicle cells, using the somatic driver *traffic jam-Gal4* (*tj-Gal4*). *hemK2*-STKD females became sterile with rudimentary ovaries (Fig. S1B, C). These ovaries often presented with fused egg chambers and disorganized follicle cells (Fig. S1C). Remarkably, the fertility and developmental disruptions were nearly fully rescued by expressing the *hemK2* transgene with *tj-Gal4* (Fig. S1B, C), suggesting that *hemK2* is crucial in both germline and somatic cell functions during oogenesis.

### Pleiotropic phenotypes in *hemK2*-GLKD ovaries indicate essential roles in oogenesis

Ovaries with *hemK2*-GLKD exhibited multiple phenotypic aberrations: hyperplasia of the posterior follicle cells, chromosomal dispersion failure, and extensive apoptosis in germline cells (Fig. 2). All egg chambers in the control females were encapsulated by a monolayer of follicle cells, whereas the majority of egg chambers in *hemK2*-GLKD females demonstrated multi-layered follicle cells in the posterior region (Fig. 2A; 94%). In addition, an increased presence of mitotic cells, marked by phospho-histone H3 (pH3), was observed in the hyperplastic multi-layered follicle cells of *hemK2*-GLKD during stages 7/8 (Fig. 2A).

**Figure 2:**
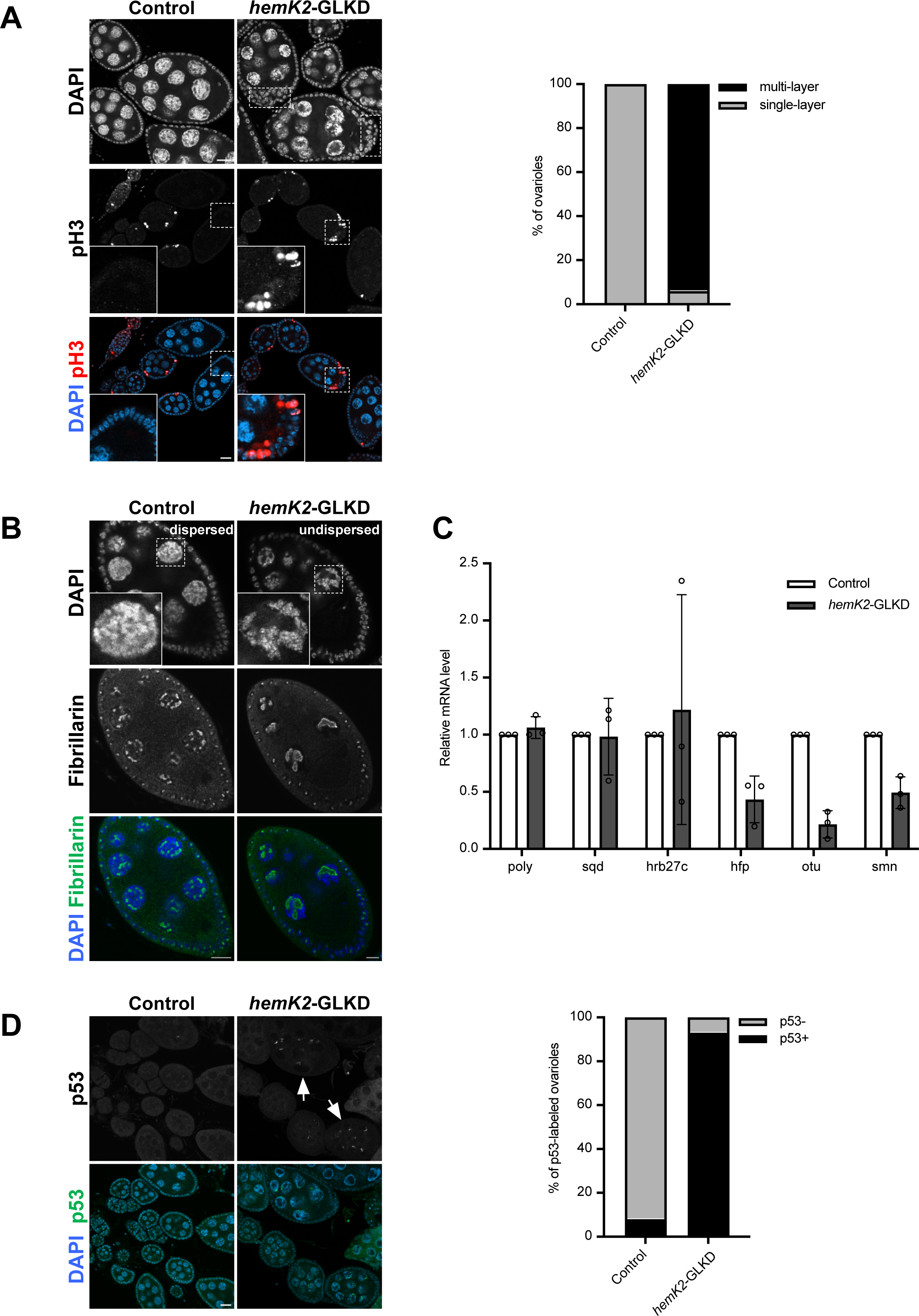
Knockdown of hemK2 leads pleiotropic defects in germline development. (A) Ovaries displaying mitotic marker phospho-histone H3 (pH3) (red) and DAPI (blue) (left panel). Dotted rectangles highlight hyperplastic posterior follicle cells in *hemK2*-GLKD (upper right panel). Scale bars, 20 µm. Quantification of ovarioles with single versus multi-layered follicle cells is provided (right panel, n=100). (B) Immunostaining for nucleolar marker Fibrillarin and DNA (DAPI) in stage 7/8 egg chambers. Insets show enlarged views of nurse cell nuclei with either dispersed or undispersed chromosomes. Scale bars, 10 µm. (C) Quantitative RT-PCR analysis of gene expression involved in chromosomal dispersion, with standard deviation indicated by error bars (n=3). (D) Immunostaining for apoptosis marker p53 and DNA (DAPI) in ovaries (left panel). Arrows indicate p53 signal accumulation in nurse cell nuclei. Scale bars, 20 µm. Percentage of ovarioles displaying p53 signals is quantified (right panel, n=100).

Moreover, nurse cells in *hemK2*-GLKD displayed defective chromosomal dispersion (Fig. 2B). Typically, wild-type nurse cells underwent a distinct polytene chromosome condensation into a ‘5-blob’ structure, which was fully dispersed by stage 6 (Dej and Spradling, 1999). In stark contrast, almost all *hemK2*-GLKD ovarioles contained nurse cell nuclei with chromosomal configurations retained in a blob-like state beyond this stage (Fig. 2B, Fig. S2A; 99%). To investigate the possible causes of this chromosomal defect, we examined the expression levels of genes known to be involved in chromosomal dispersion (Goodrich et al., 2004; Klusza et al., 2013). qRT-PCR analysis showed that the mRNA level of *otu* was significantly reduced in *hemK2*-GLKD ovaries (Fig. 2C), suggesting a correlation between *otu* expression and the chromosomal dispersion defect. The expression of N-terminal FLAG-tagged Otu104 isoform in *hemK2*-GLKD ovaries substantially recovered the dispersion of chromosomes (Fig. S2A), suggesting that *hemK2* supports chromosomal dispersion during oogenesis, partly by ensuring *otu* expression.

Additionally, *hemK2*-GLKD led to pronounced degeneration of egg chambers during mid-oogenesis, a process that is typically regulated by germline cell viability in response to physiological and environmental cues (Drummond-Barbosa and Spradling, 2001; McCall, 2004). Immunostaining for p53, an apoptotic inducer, revealed a marked increase in p53 signal in the nurse cell nuclei of most *hemK2*-GLKD ovarioles (Fig. 2D; 93%), suggesting cell death in mid-oogenesis. Consistently, the simultaneous knockdown of *p53* and *hemK2* significantly restored the progression of oogenesis in *hemK2*-GLKD ovaries (Fig. S2B). Furthermore, expression of p35, an inhibitor of caspases DrICE and Dcp-1 (Fuchs and Steller, 2015; Werz et al., 2005) also suppressed the cell death in *hemK2*-GLKD ovaries (Fig. S2B). While females with either *p53* knockdown or ectopic p35 expression in *hemK2*-GLKD did not fully recover egg-laying to control levels, the hatching rates of the resulting eggs were comparable to that of controls (Fig. S2C). Collectively, these data suggest that p53-dependent apoptosis occurs during mid-oogenesis in *hemK2*-GLKD, although the precise initiating mechanisms remain to be determined. Notably, chromosomal dispersion remained impaired in *hemK2*-GLKD ovaries even after cell death suppression (Fig. S2B), indicating that apoptosis and chromosomal dispersion defects are mediated by distinct pathways affected by *hemK2* downregulation.

### eRF1 methylation by HemK2’s catalytic activity is required for oogenesis

Previous studies have identified HemK2 as a methyltransferase acting on three different substrates: DNA N6-adenine (6mA) in human cultured cells (Xiao et al., 2018), histone H4 lysine 12 (H4K12) in A549 lung adenocarcinoma cells (Metzger et al., 2019), and a glutamine residue within the GGQ motif of eRF1 in yeast (Heurgué-Hamard et al., 2005) and human (Figaro et al., 2008). Nonetheless, the genuine substrate and functional role of HemK2 in multicellular organisms have remained elusive. First, we investigated the 6mA modification in genomic germline DNA of *hemK2*-GLKD through immunostaining with an anti-6mA antibody. No alteration in the pattern or signal intensity of 6mA staining was observed in *hemK2*-GLKD germline cells (Fig. S3A), suggesting that HemK2 does not mediate the methylation of genomic N6-adenine in the *Drosophila* germline. Moreover, levels of methylated H4K12 were similar between *hemK2*-GLKD and control ovaries (Fig. S3B), thereby excluding H4K12 as a primary substrate in the *Drosophila* germline.

Subsequently, we evaluated the methylation status of endogenous eRF1 in *hemK2*-GLKD ovaries. Due to the ineffectiveness of the anti-methyl-eRF1 antibody in Western blot analyses, we performed eRF1 immunoprecipitation followed by immunoblotting with an antibody against N5-methyl-glutamine (methyl-Q). The eRF1 methylation in *hemK2*-GLKD ovaries was significantly reduced to 35% of control, likely reflecting the contribution from unaffected somatic cells (Fig. 3A). Immunostaining with the anti-methyl-eRF1 antibody showed a substantial reduction of eRF1 methylation specifically in germline cells of *hemK2*-GLKD ovaries, while the expression level of eRF1 remained consistent across cell types (Fig. 3B). This finding was corroborated by the immunoprecipitation of FLAG-tagged eRF1 expressed in the germline, which exhibited minimal methylation in *hemK2*-GLKD ovaries compared to control (Fig. 3C), confirming the role of *Drosophila* HemK2 in eRF1 methylation. It is well documented that a glutamine residue within the GGQ motif of eRF1, crucial for peptide release, is consistently methylated by HemK2 across various species (Dinçbas-Renqvist et al., 2000; Figaro et al., 2008; Gao et al., 2020; Kusevic et al., 2016; Svidritskiy and Korostelev, 2018). To determine if the GGQ motif of eRF1 is the methylation target in *Drosophila*, we mutated the corresponding glutamine (Q185) to alanine (eRF1 Q185A) and examined their methylation in S2 cells (Fig. 3D). Consistent with observations in other systems, methylation of the eRF1 Q185A mutant was nearly undetectable, irrespective of *hemK2* knockdown, indicating that this mutation disrupts eRF1 methylation (Fig. 3D). Collectively, these results suggest that Q185 is the predominant methylation site of eRF1 by *Drosophila* HemK2.

**Figure 3:**
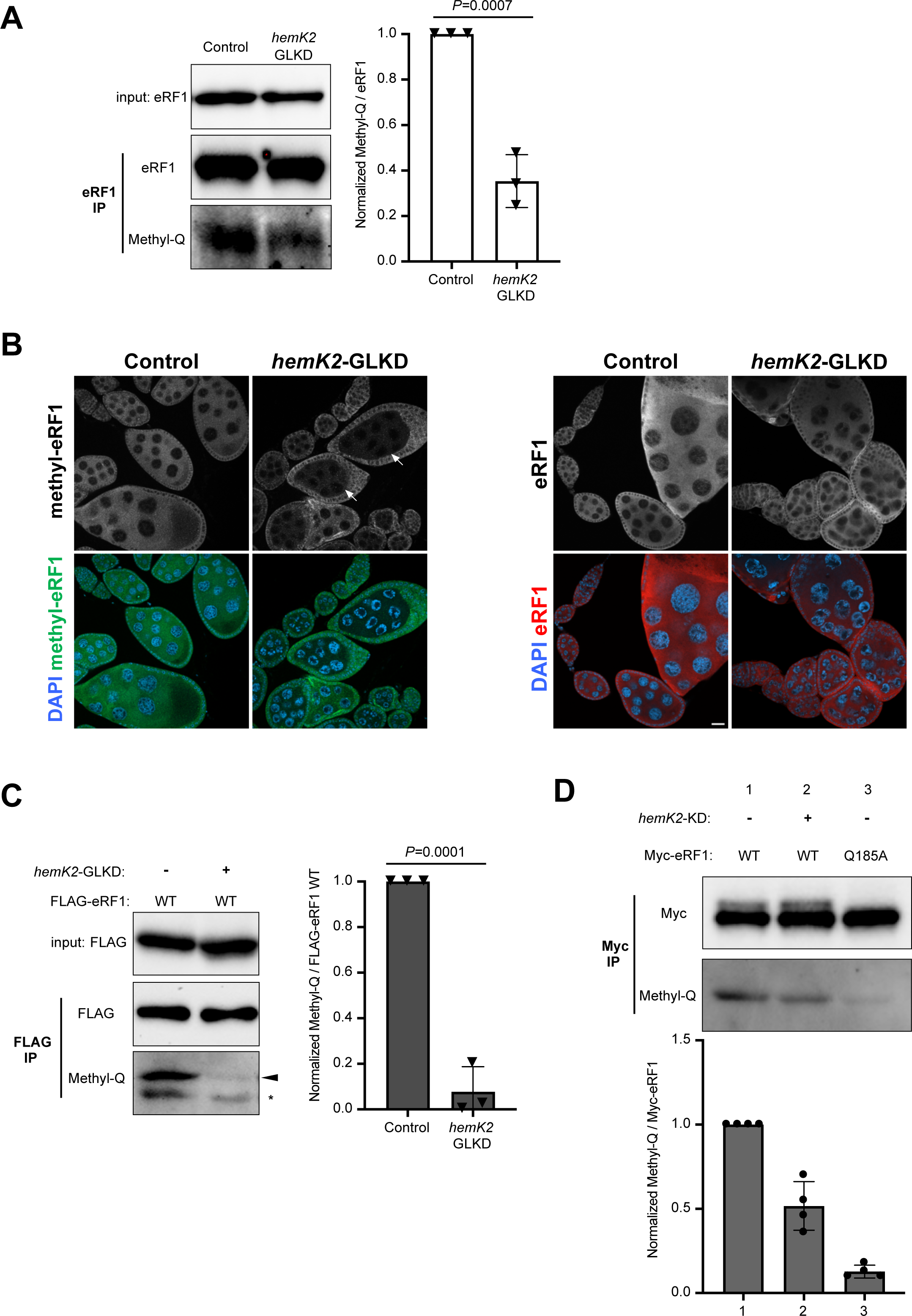
HemK2 functions for eRF1 methylation in *Drosophila*. (A) Immunoprecipitation of endogenous eRF1 and subsequent Western blot analysis for N5-methyl-glutamine (Methyl-Q) from ovarian lysates, indicating a significant reduction in eRF1 methylation in *hemK2*-GLKD compared to control ovaries (bottom panel). Quantification of methyl-Q band intensity normalized to eRF1 signal is shown (right panel), with error bars representing standard deviation (n=3) and *p*-value indicated (unpaired *t*-test). (B) Immunostaining of methylated eRF1 (green) and total eRF1 (red) in control and *hemK2*-GLKD ovaries. Arrows indicate the cytoplasm of germline cells. Scale bars, 20 µm. (C) Immunoprecipitation of germline-expressed FLAG-tagged eRF1 from ovaries. Western blot for methyl-Q illustrates a notable decrease in methylation of FLAG-eRF1 in *hemK2*-GLKD. Nonspecific bands are marked with asterisks. Right panel shows quantification of methyl-Q band intensity normalized to FLAG-eRF1, as in (A), with error bars denoting standard deviation (n=3) and *p*-value provided (unpaired *t*-test). (D) Immunoprecipitation of Myc-tagged eRF1 (WT or Q185A mutant) in S2 cells, with Western blot detection for methyl-Q. Quantifications of methyl-Q band intensities normalized to Myc-eRF1 are shown with error bars representing standard deviation (n=3).

The abovementioned rescue experiments validated HemK2’s indispensable role *in vivo* for egg development (Fig. 1 and Fig. S1). We proceeded to evaluate the significance of amino acid residues critical for HemK2’s methyltransferase activity during oogenesis. The NPPY motif of HemK2, highly conserved and recognized as a substrate-binding pocket, is vital for its methylation function in human cell lines (Fig. 4A) (Gao et al., 2020; Metzger et al., 2019). It has been reported that mutations in residues proximal to the substrate, specifically Asn122 and Tyr125 within the NPPY motif of human HemK2, abolish its catalytic activity for eRF1 methylation *in vitro* (Fig. 4B) (Metzger et al., 2019). Correspondingly, we introduced mutations at analogous positions in *Drosophila* HemK2, Asn116 and Tyr119, substituting them with alanine (N116A and Y119A, respectively), and expressed these mutants in the germline of *hemK2*-GLKD ovaries (Fig. 4C; Fig. S4A). The Y119A mutant did not recover the observed defects, such as egg chamber degeneration, egg laying and hatching rate, however, the N116A mutant restored these parameters to levels comparable to the control (Fig. 4C, D). These findings suggest that Tyr119, as opposed to Asn116, is critical for HemK2’s function within *Drosophila* germline cells.

**Figure 4:**
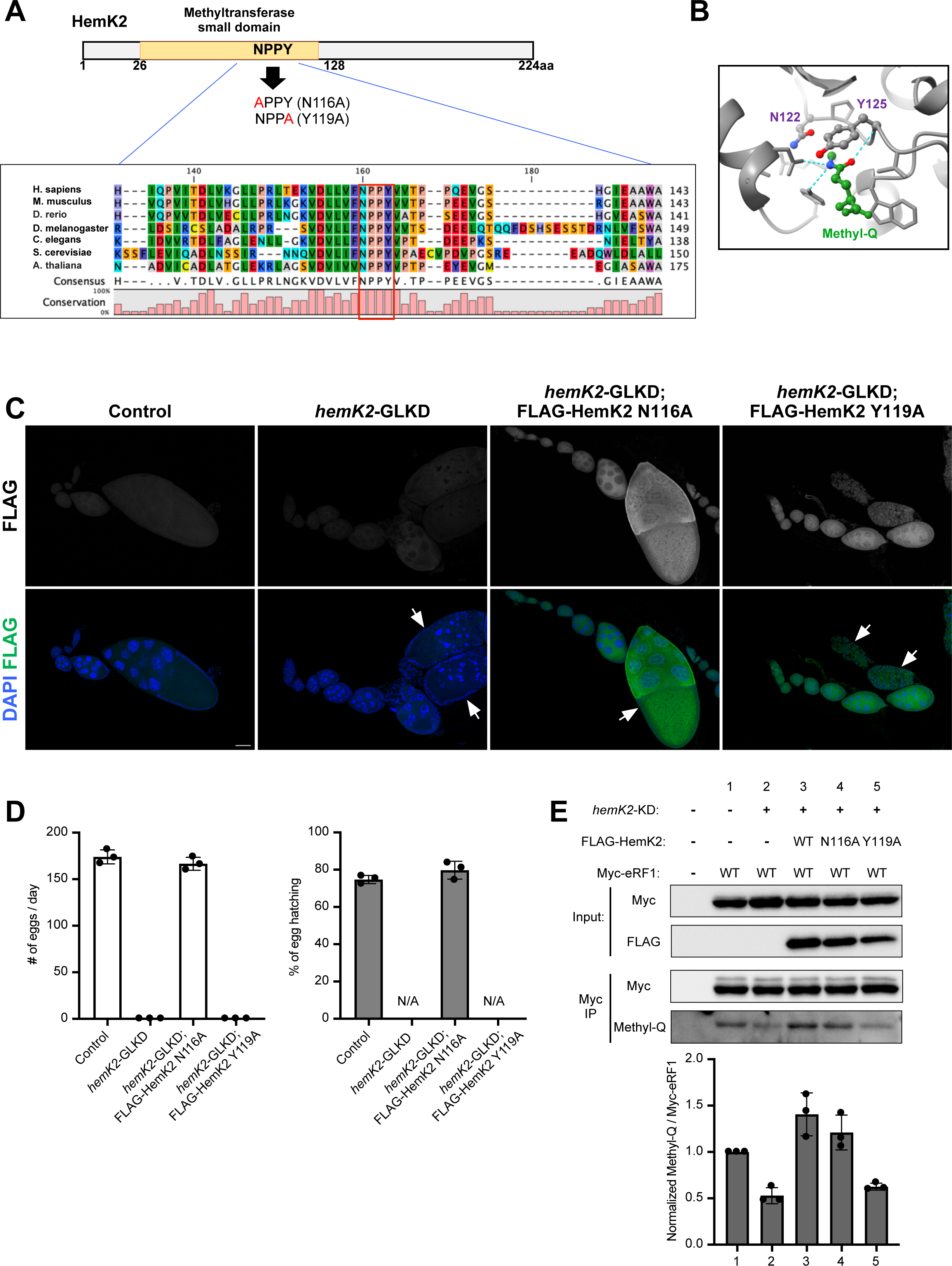
The conserved NPPY motif of HemK2 is critical for eRF1 methylation. (A) Illustration of the methyltransferase (MTase) domain within *Drosophila* HemK2, highlighting the highly conserved NPPY motif across various species. Asn116 and Tyr119 of *Drosophila* HemK2 are mutated to Ala, generating N116A and Y119A, respectively. (B) Crystal structure depiction of human HemK2-TRMT112 complexed with methyl-glutamine (Methyl-Q) and S-adenosylhomocysteine (SAH) (Gao et al., 2020), focusing on the active site proximal to methyl-Q. HemK2 is depicted in gray, methyl-Q in green with hydrogen bonds around the methyl-Q. N122 and Y125 of human HemK2 correspond to N116 and Y119 in *Drosophila* HemK2, respectively. (C) Immunostaining for FLAG-tagged HemK2 variants against a DNA counterstain (DAPI) in ovaries of the respective genotypes. Perturbation of proper oogenesis in *hemK2*-GLKD ovaries is restored by the expression of the N116A HemK2 variant but not the Y119A. Scale bars, 50 µm. (D) Analysis of egg laying and hatching rate for each genotype as described in (C). The number of eggs were counted daily for groups of three females (n=3), with standard deviation shown as error bars. (E) Immunoprecipitation followed by Western blot analysis for methyl-Q in S2 cells. Co-transfection of FLAG-tagged HemK2 and Myc-tagged eRF1, with or without *hemK2* knockdown (KD), demonstrates that eRF1 methylation loss in *hemK2*-KD is restored by wildtype HemK2 and the N116A mutant but not by the Y119A mutant. The lower panel provides quantification of methyl-Q intensity relative to Myc-eRF1 with standard deviation (n=3).

Furthermore, we investigated HemK2’s activity in catalyzing the methylation of the glutamine residue within the GGQ motif of eRF1 in S2 cells. After co-transfecting each FLAG-tagged HemK2 variant with Myc-tagged eRF1 into S2 cells and knocking down endogenous *hemK2*, we assessed eRF1 methylation (Fig. 4E). Immunoprecipitation of Myc-eRF1 followed by immunoblotting with an anti-methyl-Q antibody showed significant reduction in eRF1 methylation upon endogenous *hemK2* knockdown. This reduction was rescued by expressing wildtype HemK2 and the N116A mutant, but not by the Y119A mutant (Fig. 4E). This demonstrates that Tyr119, but not Asn116, is indispensable for the methylation of eRF1 at the 185th glutamine by *Drosophila* HemK2.

Additionally, given that HemK1, a paralog of HemK2 with a conserved NPPY motif and 42% sequence homology, is known to methylate the glutamine in the GGQ motif of mitochondrial release factors in human cells (Fang et al., 2022; Ishizawa et al., 2008), we sought to determine the specificity of cytosolic eRF1 methylation by HemK2. In S2 cells, where *hemK2* knockdown reduced eRF1 methylation, expression of HemK2, but not HemK1, rescued the methylation levels, signifying that cytosolic eRF1 methylation is an exclusive function of HemK2 (Fig. S4B).

### HemK2-mediated eRF1 methylation is required for the efficient translation and mRNA stability

We investigated the role of eRF1 methylation in oogenesis by knocking down *eRF1* expression in germline cells. *eRF1* germline knockdown (*eRF1*-GLKD) led to rudimentary ovaries that failed to produce eggs, a phenotype more severe than that of *hemK2*-GLKD (Fig. 1B, 5A, B). Immunostaining with Vasa (Vas), a DEAD box RNA helicase and germline marker, indicated agametic germaria with a stark reduction in germline cells in *eRF1*-GLKD ovaries (Fig. 5C). The defects in *eRF1*-GLKD were successfully rescued by expressing FLAG-tagged wildtype eRF1, but not with the methylation-deficient Q185A mutant in germline cells (Fig. 5B, C). This highlights the critical role of the 185^th^ glutamine residue, and likely its methylation, in germline cell viability and oogenesis progression. The less severe phenotype in *hemK2*-GLKD could result from incomplete disruption of eRF1 methylation, potentially due to residual *hemK2* expression or compensation by other methyltransferase(s).

**Figure 5:**
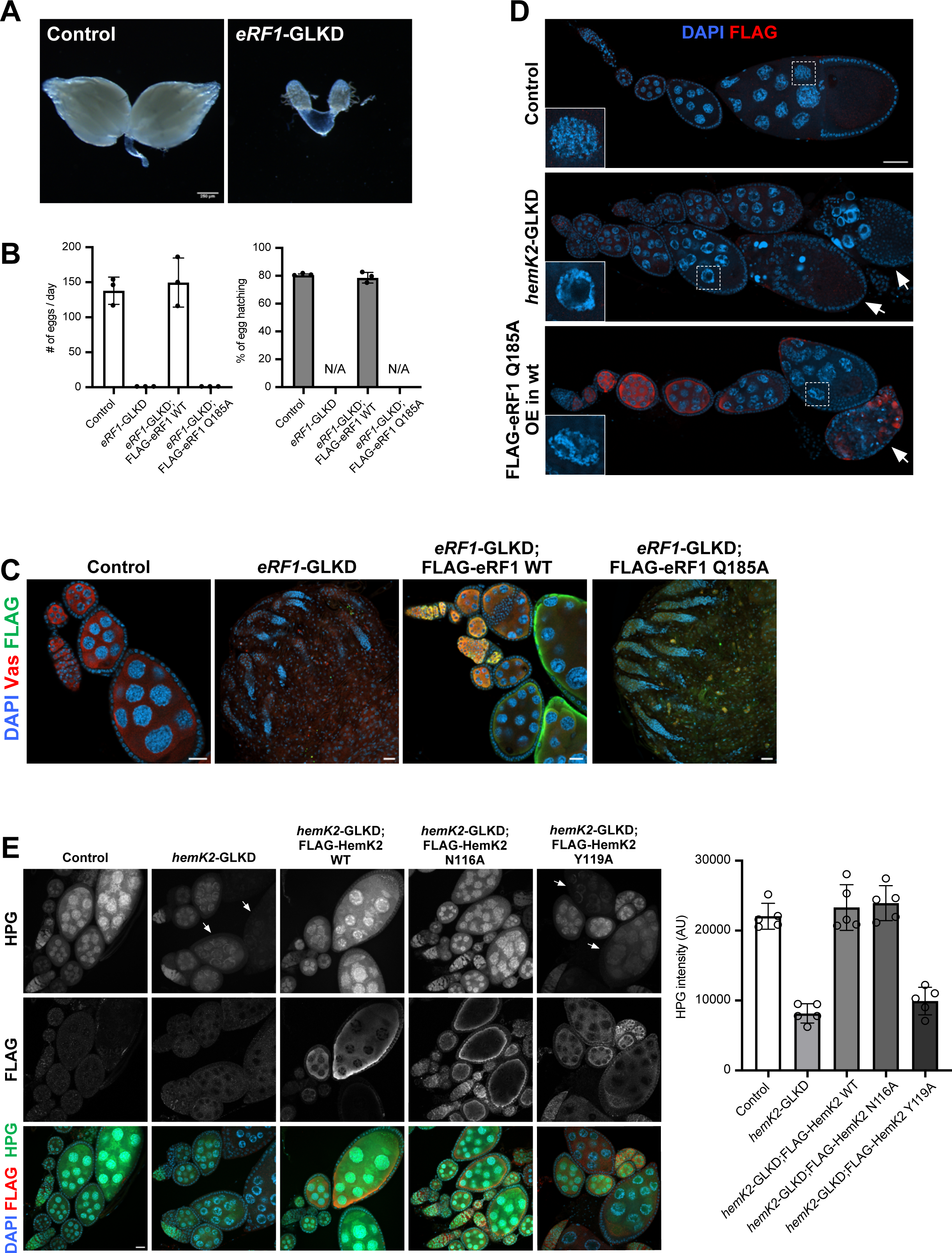
eRF1 methylation ensures proper progression of oogenesis and efficient translation. (A) Comparative morphological images of control and *eRF1* germline knockdown (*eRF1*-GLKD) ovaries. Scale bar, 250 µm. (B) Analysis of egg production and hatching rate across different genotypes. The numbers of the eggs were counted daily for groups of three females (n=3), with error bars representing standard deviation. (C) Immunostaining of wild-type and Q185A mutant FLAG-tagged eRF1 (green), with Vas (red) marking germline cells, and DAPI (blue) for DNA, in ovaries corresponding to each genotype listed in (B). Loss of germline cells by *eRF1*-GLKD is restored with wild-type eRF1 but not with the Q185A mutant. Scale bars, 20 µm. (D) Immunostaining for FLAG-tagged eRF1 Q185A (red) and DNA (DAPI, blue) in ovaries. Overexpression of FLAG-eRF1 Q185A in wild-type background recapitulates the phenotypic consequences of *hemK2*-GLKD. Arrows point to degenerating egg chambers. Insets show enlarged views of nurse cell nuclei. Scale bars, 50 µm. (E) Protein synthesis assay using L-homopropargylglycine (HPG) in ovaries for each genotype indicated (green in left panels), with concurrent immunostaining for FLAG-HemK2 (red). Scale bars, 20 µm. The right panel shows quantification of HPG signal intensity, with measurements from representative images (n=5) and standard deviation as error bars.

Moreover, expressing FLAG-eRF1 Q185A mutant in germline cells of wildtype ovaries disrupted egg production (Fig. 5D). This expression led to developmental arrest at mid-oogenesis, with defects in chromosome dispersion and extensive apoptosis, mirroring the *hemK2*-GLKD phenotype (Fig. 5D; Fig. S5A, B). Over 80% of ovarioles expressing eRF1 Q185A exhibited p53-positive egg chambers (Fig. S5C). Therefore, the expression of eRF1 Q185A effectively recapitulates the *hemK2*-GLKD phenotype, supporting that the absence of eRF1 methylation primarily causes the observed defects.

Given that eRF1 GGQ motif methylation is essential for nascent peptide release and translational termination efficiency, and that its perturbation could lead to translational deficits by hindering ribosome recycling, we assessed translation activity in *hemK2*-GLKD ovaries using an HPG incorporation assay *ex vivo*. HPG, a methionine analog with an alkyne group, can be incorporated into newly synthesized proteins and is visualized through a ‘Click reaction’ with a fluorescent dye (Shen et al., 2021). A notable decrease in HPG signal in cytoplasm of germline cell was observed in *hemK2*-GLKD ovaries (Fig. 5E), indicating a requirement for eRF1 methylation in maintaining translation efficiency. This translational reduction in *hemK2*-GLKD was completely restored by expressing either wildtype HemK2 or the N116A mutant, but not by the Y119A mutant (Fig. 5E). Similarly, germline expression of the methylation-deficient eRF1 Q185A mutant abrogated translation activity (Fig. S5D). Collectively, these results underscore the necessity of HemK2-mediated eRF1 methylation for efficient translation in the *Drosophila* ovary.

To elucidate the molecular deficiencies resulting from *hemK2*-GLKD and eRF1 dysfunction, we conducted ovarian mRNA-seq analysis using next-generation sequencing technology. Late egg chambers in control ovaries were selectively removed to minimize developmental discrepancies between *hemK2*-GLKD and control samples. Aligning with our RT-PCR data (Fig. 1C and Fig. 2C), mRNA-seq confirmed the marked downregulation of *hemK2* (log2FC=-1.27) and *otu* (log2FC=-1.45) (Supplementary Table S4). Additionally, we identified 478 upregulated and 325 downregulated genes (log_2_FC < −1 or > 1; q-val < 0.01). Gene Ontology (GO) term analysis (Thomas et al., 2022) of differentially expressed genes indicated a significant downregulation in genes associated with female reproductive processes, such as female gamete generation, oogenesis, and germ cell development (Fig. 6A), while genes unrelated to oogenesis were comparatively upregulated (Fig. 6B).

**Figure 6:**
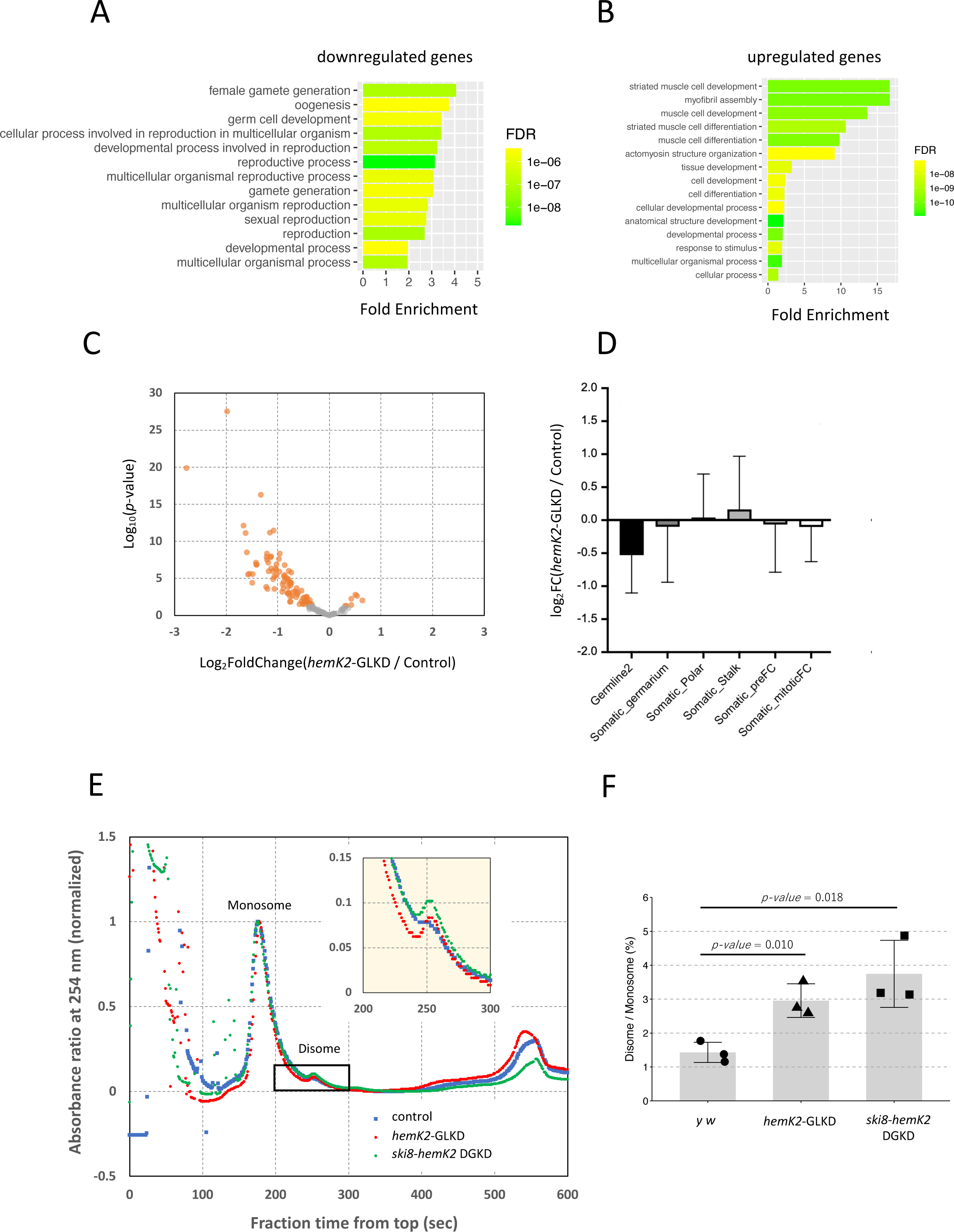
Knockdown of *hemK2* leads mRNA reduction and ribosome stalling. (A) Gene Ontology (GO) analysis of genes downregulated in *hemK2*-GLKD ovaries. Enrichment of GO biological process terms is analyzed by PANTHER (version 18.0). The fold enrichments are plotted and colored by their False Discovery Rate (FDR) values. (B) GO analysis for genes upregulated in *hemK2*-GLKD ovaries, conducted similarly to panel (A). (C) Volcano plot representing the differential expression of germline-enriched genes in *hemK2*-GLKD ovaries, with significance threshold set at *p*-value < 0.05 and log_2_FC < −1 or =1 (orange). Gene classification is based on single-cell mRNA-seq analysis of *Drosophila* ovaries (Jevitt et al., 2020). (D) Log_2_-fold changes of cell-type enriched genes in *hemK2*-GLKD ovaries relative to control, with mean values and standard deviation depicted. (E) Ribosome profiling in ovarian lysates from control (blue), *hemK2*-GLKD (red), and double-germline knockdown of *hemK2* and *ski8* (green). Monosome and disome were separated using sucrose gradient sedimentation. Absorbance at 254 nm was recorded to generate elution profiles, normalized to the monosome peak for each sample. Insets highlight the disome peaks. (F) Disome to monosome peak ratios were calculated across three experimental replicates. Bar graphs represent average ratios with standard deviation (n=3) and *p*-value (unpaired *t*-test).

To further dissect the gene expression changes following germline knockdown of *hemK2*, we referenced a previous study detailing gene expression across different ovarian cell types from single-cell mRNA analyses; many different cell types were distinguished based on gene expression patterns. Focusing on genes expressed in one germline and five somatic cell types representing early to mid-oogenesis, we observed a significant downregulation (average log_2_FC = −0.55) in *hemK2*-GLKD for genes enriched in germline 2 cell type (N=154), which are expressed from germarium region 2 onward (Fig. 6C, D) (Jevitt et al., 2020). In contrast, genes associated with the five somatic cell types did not show notable changes (Fig. 6D; Fig. S6A-E), suggesting that *hemK2*-GLKD specifically downregulates mRNAs in germline cells.

Previous research indicates that eRF1 methylation at the GGQ motif substantially enhances the release of nascent peptides, accelerating translation termination by 10 to 100-fold (Pierson et al., 2016). We hypothesize that impaired methylation on eRF1 could decrease the peptide release rate, leading to ribosome stalling at stop codons of actively translated mRNAs. Such stalling is known to activate mRNA surveillance process, targeting mRNAs with engaged ribosomes for degradation via the No-Go Decay (NGD) pathway (Brandman and Hegde, 2016; Doma and Parker, 2006; Ikeuchi et al., 2019; Simms et al., 2017; Tomomatsu et al., 2023). To investigate this hypothesis, we analyzed ribosome stalling in *hemK2*-GLKD by performing a sucrose gradient sedimentation assay with ovarian lysates. The lysates, treated with emetine to halt translation and RNase A to digest unprotected mRNA regions, revealed a predominance of monosomes and a negligible disome presence in the control, indicative of minimal ribosome stalling and collision (Fig. 6E). However, in *hemK2*-GLKD, disome levels increased approximately 2.1-fold (Fig. 6F, *hemK2*-GLKD), suggesting that ineffective nascent polypeptide release due to unmethylated eRF1 leads to ribosome collision.

### Blockage of the No-Go Decay pathway restores mRNA destabilization and mid-oogenesis arrest in *hemK2*-GLKD ovaries

To ascertain if the No-Go Decay (NGD) pathway was responsible for the pronounced defects observed in *hemK2*-GLKD, we conducted a simultaneous knockdown of the mRNA surveillance pathway components along with *hemK2* in germline cells (Fig. 7 and Fig. S7A). In the double-germline knockdown (DGKD) of *hemK2* and *ski8*—a component of the Ski complex that collaborates with the exosome and facilitates 3’ to 5’ mRNA degradation (Zinoviev et al., 2020)—, developmental impairments associated with *hemK2*-GLKD were partially restored (Fig. 7A). Females from the *ski8*-*hemK2* DGKD had more advanced stages of egg chambers and laid a comparable number of eggs to controls, though their hatching rate was reduced to 40% of the control (Fig. 7A, B). Considering the Ski complex’s role in general RNA decay, we further investigated additional NGD factors, such as *pelota* (*pelo*), which is crucial for dissociating stalled ribosomes and promoting RNA decay within the NGD and No-Stop Decay (NSD) pathways (Becker et al., 2011; Jamar et al., 2017; Kobayashi et al., 2010; Tsuboi et al., 2012). Remarkably, *pelo*-*hemK2* DGKD females exhibited a similar phenotype to that of the *ski8-hemK2* DGKD, with advanced egg chamber development and a 60% hatching rate, comparable to the number of eggs laid (Fig. 7A, B). Furthermore, the mid-oogenesis arrest induced by expressing the dominant-negative eRF1 Q185A mutant was also partly rescued by the germline knockdown of either *ski8* or *pelo* (Fig. S7B), reinforcing the notion that the eRF1 methylation defect triggers mRNA surveillance mechanisms.

**Figure 7:**
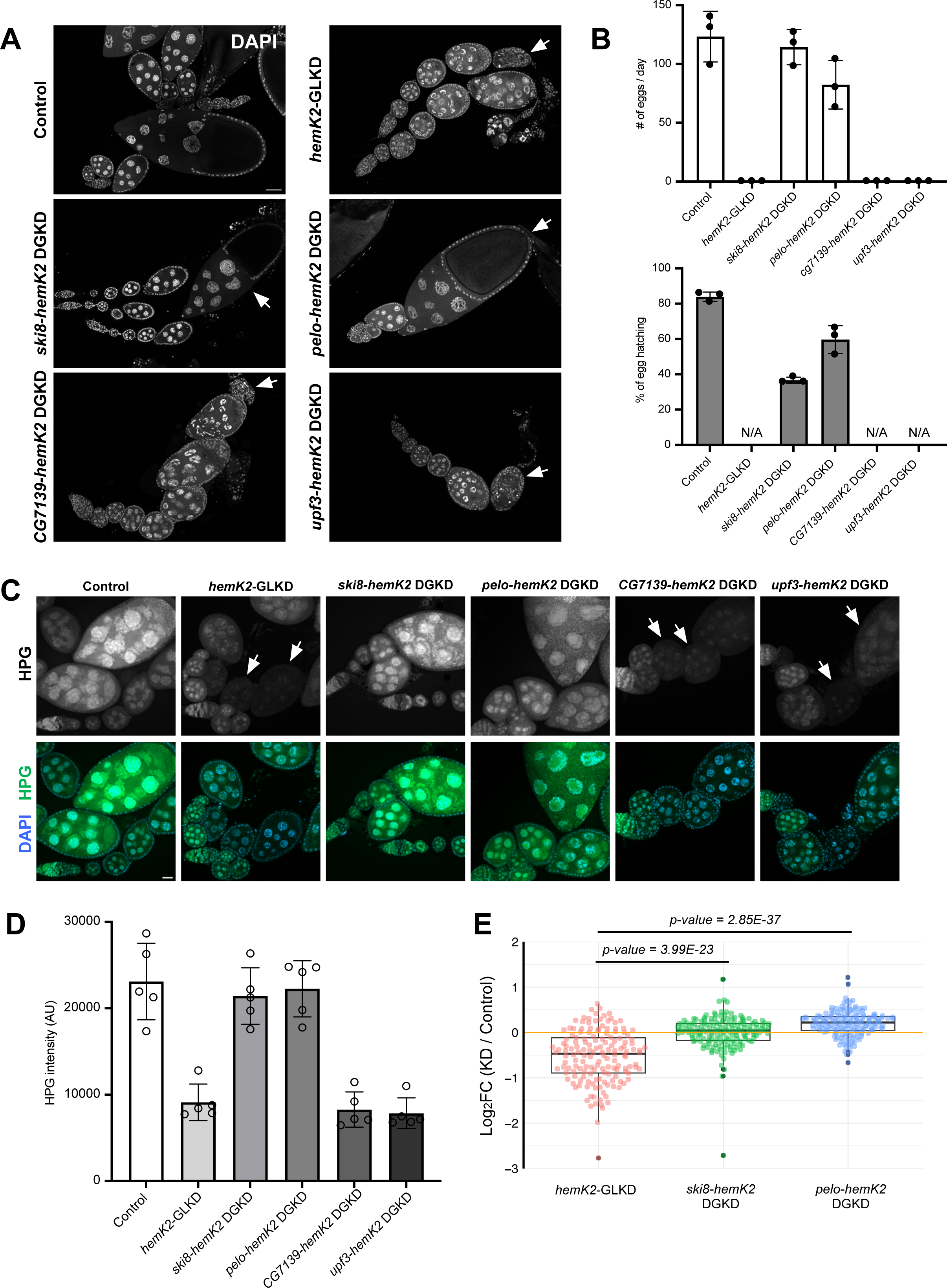
Blockage of No-Go-decay pathway restores oogenesis defects in *hemK2*-GLKD ovaries. (A) Fluorescent staining with DAPI for DNA in ovaries from various genotypes. Rescue of the oogenesis defects observed in *hemK2*-GLKD is achieved through combined knockdown of *ski8* or *pelo*, but not with *CG7139* or *upf3*. Scale bars, 50 µm. (B) Analysis of egg production and hatching rate for the indicated genotypes. The numbers of egg counted for groups of three females each day (n=3), with error bars representing standard deviation. (C) Assay of protein synthesis in ovaries using L-homopropargylglycine (HPG), across the specified genotypes. Scale bars, 20 µm. (D) Quantitative analysis of HPG fluorescence intensity for each genotype, with measurements taken from representative images (n=5). Error bars denote standard deviation. (E) Box plot depicting the log_2_-fold changes of genes enriched in germline 2 cell type in ovaries from *hemK2*-GLKD, *ski8*-*hemK2* DGKD, and *pelo-hemK2* DGKD compared to control. Paired *t*-test *p*-values are presented, comparing *hemK2* single knockdown with each respective double knockdown.

Investigating the role of the NGD pathway in the defects seen in *hemK2*-GLKD ovaries, we assessed another putative NGD component, *nonu-1*/*cue2*, an endonuclease identified in *C. elegans* and *S. cerevisiae* that cleaves mRNA near a stalled ribosome during translation (D’Orazio et al., 2019; Glover et al., 2020) (Fig. S7C). Contrasting with the partial rescue observed with *ski8* and *pelo* knockdowns, the simultaneous germline knockdown of *CG7139*, encoding the *Drosophila* ortholog of *nonu-1*/*cue2*, and *hemK2* did not ameliorate the observed phenotypic defects (Fig. 7A, B; Fig. S7A). This indicates that *CG7139* may not be an essential component of the NGD pathway in *Drosophila*, or it may act redundantly with other endonucleases in the translation-coupled mRNA surveillance mechanism compromised by *hemK2* knockdown. Further analysis revealed that a dual germline knockdown of *hemK2* and *upf3*, a factor in Nonsense-Mediated Decay (NMD) (Tan et al., 2022), did not reverse the phenotypic abnormalities in *hemK2*-GLKD ovaries (Fig. 7A, B), emphasizing a specific role of NGD in mediating mRNA degradation. The efficiency of the knockdown for each component was verified by qRT-PCR (Fig. S7A; ranging from 10 to 40% relative to control), accounting for the contribution of non-knockdown somatic cells to the total ovarian RNA, confirming a significant reduction in expression levels. Consistent with their roles in rescuing developmental morphology, the HPG incorporation assay demonstrated that knockdown of *ski8* or *pelo*, but not *CG7139* or *upf3*, in *hemK2*-GLKD ovaries partially restored translational activity (Fig. 7C, D). These results support the hypothesis that disrupted eRF1 methylation in *hemK2*-GLKD leads to compromised translation termination and ribosome stalling, specifically triggering mRNA degradation via the NGD pathway.

In alignment with the recovery of oogenesis defects in *hemK2*-GLKD, the expression levels of downregulated protein-coding RNAs were restored by the additional germline knockdown of either *ski8* or *pelo* (Fig. 7E, Fig. S6F, G), further validating the specific activation of NGD upon *hemK2*-GLKD. The concurrent germline knockdown of *ski8* or *pelo* with *hemK2* also reinstated general protein synthesis, despite the observation of ribosome stalling to a similar degree as in the single *hemK2* knockdown (Fig. 6E, F). These findings suggest that when mRNAs are not eliminated by NGD, translation re-initiation may occur, enabling protein synthesis and partially recovering egg maturation (Fig. 7A, B). Overall, our results indicate that perturbations in eRF1 methylation due to *hemK2*-GLKD result in the inefficient release of synthesized peptides and the activation of the NGD pathway, likely due to ribosome stalling.

## Discussion

The reduction of *hemK2* expression led to female sterility, marked by a significant interruption of oogenesis in both germline and somatic cells of the ovary, culminating in developmental arrest during the mid-stages (Fig. 1; Fig. S1). Notably, *hemK2*-GLKD ovaries exhibited morphological aberrations, such as follicle cell hyperplasia and chromosomal dispersion failure (Fig. 2A, B). It has been previously reported that the expression of the Delta ligand in germline cells activates Notch-Delta signaling pathway, which prompts the transition from the mitotic cell cycle to the endocycle in follicle cells (Deng et al., 2001; Schaeffer et al., 2004; St. Johnston, 2001). The hyperplasia observed in the posterior follicle cells of *hemK2*-GLKD ovaries might be due to diminished Delta ligand expression in germline cells, leading to an extended period of mitotic division in follicle cells beyond the stage 6.

In egg chambers of stage 6 and beyond, *hemK2*-GLKD ovarioles predominantly contained nurse cell nuclei with unspread chromosomes (Fig. 2B; Fig. S2A). Inhibition of cell death in *hemK2*-GLKD ovaries significantly improved egg chamber formation, yet the chromosomal dispersion defect persisted (Fig. S2B). These findings imply that germline cell death in *hemK2*-GLKD is not caused by the failure of chromosomal dispersion but likely due to ribosome stalling, which is a consequence of eRF1 methylation disturbance (Wu et al., 2020). It has been reported that ribosome collisions can initiate two distinct stress response pathways: a transient collision may trigger a GCN2-mediated pathway promoting cell survival, while more severe collisions could lead to the activation of MAPKK, inducing apoptosis in human cell lines (Wu et al., 2020). Our data suggest that perturbation of methylation due to *hemK2*-knockdown results in ribosome stalling and subsequent collisions, triggering apoptosis in the translationally active environment of *Drosophila* ovaries.

Although HemK2 in yeast and mammalian cells has been shown to mediate methylation of DNA 6mA, histone H4K12, and eRF1 (Figaro et al., 2008; Metzger et al., 2019; Xiao et al., 2018), our findings indicate that in *Drosophila*, HemK2 does not affect the methylation of DNA 6mA or histone H4K12 (Fig. S3A, B). Instead, *hemK2* knockdown led to a substantial decrease in eRF1 methylation (Fig. 3A-C), underscoring eRF1 as the specific substrate for *Drosophila* HemK2. Notably, Tyr119, but not Asn116, is essential for HemK2’s methyltransferase activity, which is pivotal for oogenesis and eRF1 methylation in *Drosophila* (Fig. 4C-E). This contrasts with the necessity of both corresponding residues in human HemK2 for eRF1 methylation in vitro (Metzger et al., 2019). However, recent structural analyses of human HemK2 revealed that the Asn122 side chain does not interact with the methylated glutamine within the substrate binding pocket (Gao et al., 2020), aligning with the dispensability of Asn116 in *Drosophila* HemK2’s methyltransferase activity.

Murine HemK2 substrate specificity studies using a peptide array have delineated a putative consensus motif: G-Q-[SRYKLGAMTC]-[ARFGLWYCSQKH]-[LARCQFYT]-R, where square brackets indicate any of the residues listed therein (Kusevic et al., 2016). Employing this motif, we have identified several *Drosophila* proteins, including CG17841, Spargel, Chd1, Ythdc1, and eRF1, as potential HemK2 substrates. These candidates may undergo methylation by HemK2 under certain conditions. However, the phenotypes associated with the overexpression of the methylation-deficient eRF1 Q185A mutant mirrored those observed in *hemK2*-GLKD (Fig. 5D), proposing that in the context of *Drosophila* germline cells, eRF1 is the primary substrate for HemK2.

Consistent with previous reports, methylation of the conserved GGQ motif within release factors has been shown to significantly enhance nascent peptide release, accelerating the termination rate of protein synthesis by 10 to 100-fold (Pierson et al., 2016). Corroborating this, our study found that perturbation of eRF1 methylation by *hemK2*-GLKD led to a marked decrease in translation efficiency (Fig. 5E). Notably, the reduced protein synthesis, alongside the defects in oogenesis and the downregulation of mRNAs, were substantially rescued by inhibiting general RNA degradation and NGD pathway, while NMD blockade did not yield similar results (Fig. 7A-D). These findings imply that the ribosome, even when associated with unmethylated eRF1, can synthesize minimal yet essential polypeptides, provided that the mRNAs are available for translation.

Our findings support the role of HemK2 as an authentic protein methyltransferase in *Drosophila*, crucially involved in modulating eRF1 activity via glutamine methylation, a modification vital for appropriate germline development. Germline nurse cells efficiently produce proteins and mRNAs destined for transfer to the oocyte, necessitating stringent regulation of resource-intensive protein synthesis. Nonetheless, the specific regulatory mechanisms governing eRF1 methylation by HemK2 remain elusive and warrant additional exploration. It is plausible that this mechanism serves to optimize egg production under favorable physiological conditions, while downregulating it when conditions are less than ideal.

## Materials and Methods

### Fly stocks and cultures

All *Drosophila* stocks and crosses were raised at room temperature or at 25°C on standard food (5% (w/v) dry yeast, 5% (w/v) corn flower, 2% (w/v) rice bran, 10% (w/v) glucose, 0.7% (w/v) agar, 0.2% (v/v) propionic acid, and 0.05% (w/v) *p*-hydroxy butyl benzoic acid). For RNAi-mediated knockdown experiments, the following transgenic RNAi Project (TRiP) lines from the Bloomington Drosophila Stock Center (BDSC, USA), were utilized: *hemK2* (HMS02003, #40837), *eRF1* (HMS05812, #67900), *upf3* (HMJ22158, #58181), *cue2* (HMS02721, #44007), *ski8* (HMC04664, #57377), and *p53* (HMS02286, #41720). The attP40 (#25709) and attP2 (#25710) lines were employed for transgene injections. The following drivers were used to express transgenes or shRNAs: *NGT40* and *nosGal4-VP16* (Grieder et al., 2000), *Matα-tub-Gal4* (#7063), and *traffic jam-Gal4* (Drosophila Genetic Resource Center, Japan, #104055) (Hayashi et al., 2002). The UASp-P35 transgenic line was generously provided by Dr. Bergmann (Werz et al., 2005). Controls for each experiment were either the *y w* strain or the corresponding heterozygous line.

### Generation of transgenic fly lines

The transgenic fly lines generated were as follows: FLAG-HemK2 wildtype at attP2, FLAG-HemK2 N116A at attP2, FLAG-HemK2 Y119A at attP2, FLAG-Snapin at attP2, FLAG-eRF1 wildtype at attP2, FLAG-eRF1 Q185A at attP2, and FLAG-Otu104 at attP2. In each instance, the coding sequence or its mutant variant was amplified by PCR, utilizing *y w* genomic DNA or cDNA synthesized from *y w* ovarian RNA as templates. A triple FLAG tag sequence was appended to the N-terminal region of the amplification products. The purified PCR fragments were subsequently cloned into the *Xba*I-linearized pUASp-K10-attB vector (Koch et al., 2009) via In-Fusion reaction (#639648, Takara Bio, Japan). Plasmids confirmed to be correct were injected into embryos of nos-phiC31;;P{CaryP} attP2 (#25710) following standard protocols.

The *hemK2* TRiP construct at attP2 and *pelo* TRiP constructs at attP2 and attP40 sites, were also created. These TRiP constructs were produced by ligating annealed oligonucleotides (TRiP-Fw/Rv) into the *Nhe*I- and *Eco*RI-linearized pWalium22 vector (Perkins et al., 2015). UASp and Walium22 plasmids were then injected into embryos of nos-phiC31; P{CaryP}attP40 (#25709) or nos-phiC31;;P{CaryP} attP2 (#25710). The sequences of the oligonucleotides used for annealing are provided in Supplementary Table S1.

### Fluorescent immunostaining

Immunostaining was performed as previously described (Lim et al., 2022). Ovaries were dissected and slightly teased apart in phosphate-buffered saline (PBS), then fixed with 5.3% (v/v) paraformaldehyde (PFA, Electron Microscopy Sciences, USA) in PBS for 10 min at room temperature. The fixed ovaries were washed for 30 min with several changes of PBX (PBS containing 0.2% (v/v) Triton X-100). This was followed by a blocking step involving incubation in PBX containing 4% (w/v) bovine serum albumin (BSA) for a minimum of 30 min. Ovaries were incubated with primary antibodies diluted in PBX containing 0.4% (w/v) BSA overnight at 4°C, then washed for 1 h with several changes of PBX. Secondary antibodies were applied for over 2 hours at room temperature; following this, ovaries were washed for 1 h with PBX and stained with 1 μg/ml of 4′,6-diamidino-2-phenylindole (DAPI) (SIGMA, USA) for 10 min in PBX. After a final rinse with PBS, the ovaries were mounted onto slides using Fluoro-KEEPER Antifade (Nacalai Tesque, Japan)

The following primary antibodies were used in the indicated dilution; mouse anti-FLAG (SIGMA [M2], F1804, 1:200), guinea pig anti-Vas (1:2,000) (Patil and Kai, 2010), rabbit anti-pH3 (Millipore, 06-570, 1:250), mouse anti-p53 (DSHB [25F4], 1:100), rabbit anti-Fibrillarin (Abcam, ab5821, 1:200), rabbit anti-DNA 6mA (Synaptic Systems, 202003, 1:1,000), rabbit anti-eRF1 (SIGMA, E8156, 1:200), and rabbit anti-methyl-eRF1 (1:100) (Lacoux et al., 2020). Secondary antibodies were Alexa Fluor 488-, 555-conjugated goat antibodies against rabbit, mouse, and guinea pig IgG (Invitrogen, A11029, A11034, A11073, A21424, A21429, A21435) at a 1:200 dilution in PBX containing 0.4% (w/v) BSA. Ovarian images were captured using a Zeiss LSM 900 confocal microscope and processed with Zen (Zeiss) and Fiji (Schindelin et al., 2012) software.

### Protein synthesis assay

Protein synthesis was assessed using the Click-iT® L-homopropargylglycine (HPG) Alexa Fluor 488 Protein Synthesis Assay Kit (Molecular Probes, USA), which measures newly synthesized proteins through quantification of the fluorescent HPG signal. Freshly dissected ovaries were incubated in 1 mM HPG in PBS for 30 min and then fixed with 5.3% PFA in PBS for 10 min. Subsequently, the ovaries were permeabilized by washing three times in PBX for 30 minutes: the first wash utilized 3% BSA in PBX, followed by two additional washes in PBX alone. The ovaries were then incubated with a freshly prepared Click-iT® reaction cocktail for 30 min. After washing with Click-iT® reaction rinse buffer, ovaries were stained with HCS NuclearMask™ Blue Stain working solution (1:2,000 in PBX) for 30 min as per the manufacturer’s instructions. Subsequent immunostaining with antibodies and image acquisition were performed as described above.

### Quantification of HPG signal intensity for translation activity

Fluorescence intensity of HPG signals in nurse cells of egg chambers (between stages 3 to 8), was quantified to assess translation activity. Identically sized small squares were selected within the cytosolic areas, avoiding the nucleus. The fluorescence intensity for each square was measured after background subtraction, with the background defined as a blank region lacking any cells or signals. For each egg chamber, 2 to 5 of these squares were analyzed. The average fluorescence intensity across all squares within an image was calculated to represent the translation activity. This process was repeated for a total of 5 images per genotype. Image processing and analysis were conducted using Fiji software.

### Immunoprecipitation and Western blotting

For immunoprecipitation, 100 ovaries were dissected in cold PBS and homogenized using a pestle in PBS with 0.1% (v/v) Tween-20 (PBST), supplemented with a protease inhibitor cocktail (PIC, 11873580001, Roche, Switzerland). After centrifugation at 15,000 xg for 10 min at 4°C, the supernatant was collected. Protein G magnetic beads (TAMAGAWA SEIKI, TAS8848N1173) were pre-incubated with anti-FLAG antibody (SIGMA [M2], F1804) for more than 1 h at 4°C with gentle rotation before adding lysate. The antibody-conjugated beads were washed twice with cold PBST and then incubated with the lysate for 3-4 hours at 4°C. The proteins were washed three times with cold PBST and eluted from the beads in 50 µl of sample buffer at 95°C for 5 min.

For Western blotting, ovary lysates or immunoprecipitates were separated by SDS-PAGE and transferred onto a 0.45 µm Clear Trans® PVDF membrane (Wako, Japan). The membrane was blocked in PBST with 3% (w/v) skim milk (Nacalai Tesque, Japan) and incubated with primary antibodies diluted in HIKARI signal enhancer (Nacalai Tesque). The primary antibodies used included rabbit anti-eRF1 (SIGMA, E8156, 1:1,000), rabbit anti-methyl-Q (Millipore, ABS2185, 1:1,000), mouse anti-FLAG (SIGMA [M2], F1804, 1:1,000), mouse anti-c-Myc (Wako [9E10], 017-21871, 1:1,000), mouse anti-histone H4 (MAB Institute, 388-09171, 1:2,000), rabbit anti-H4K12met (PTM BIO, PTM-685RM, 1:2,000), mouse anti-Tubulin (Santa Cruz [DM1A], sc-32293, 1:3,000), rabbit anti-HA (Cell Signaling Technology [C29F4], 3724, 1:1,000). Horseradish peroxidase (HRP)-conjugated secondary antibodies used were goat anti-rabbit IgG or anti-mouse IgG (Bio-Rad, 170-6515 or 170-6516, 1:3,000). To prevent signal overlap from IgG heavy chain with eRF1, VeriBlot-HRP (Abcam, ab131366, 1:1,000) was used in Western blot analysis for immunoprecipitation of endogenous eRF1 or FLAG-eRF1. Chemiluminescent signals were detected using Chemi-Lumi One (Nacalai Tesque, 07880-70) and imaged by the ChemiDoc™ Touch MP system (Bio-Rad). Band intensities were quantified using Image Lab software (Bio-Rad) from independent experiments.

### Quantitative RT-PCR

Total RNA was extracted from ovaries using TRIzol LS Reagent (Invitrogen) strictly according to the manufacturer’s instructions. The purified RNA (1 µg) was then treated with DNase I (Invitrogen) to remove any genomic DNA contamination. This was followed by inactivation of the DNase I with EDTA. Reverse transcription to synthesize cDNA was conducted using the Superscript III First-Strand Synthesis System (Invitrogen), as per the manufacturer’s protocol. Quantitative PCR (qPCR) was performed using the KAPA SYBR Fast qPCR Master Mix (KAPA Biosystems) on the StepOnePlus Real-Time PCR System (Applied Biosystems). Relative expression levels were normalized to those of the reference gene *rp49* across three biological replicates. The primer sequences used to determine gene expression levels are listed in Supplementary Table S2.

### S2 cell experiments

*Drosophila* S2 cells were maintained at 28°C in Schneider’s medium, supplemented with 10% (v/v) fetal bovine serum (FBS) and antibiotics (penicillin and streptomycin). Knockdown of endogenous *hemK2* was achieved using double-stranded RNA (dsRNA) targeting a 186 bp region from the ATG start codon of the *snapin* gene (refer to Fig. S1). The dsRNA was synthesized using the AmpliScribe T7 High Yield Transcription Kit (Lucigen, USA). Primer sequences for the PCR amplification are provided in Supplementary Table S3.

For transfection, plasmids employed were Myc-HemK2 wildtype, Myc-HemK2 N116A, Myc-HemK2 Y119A, FLAG-eRF1 wildtype, FLAG-eRF1 Q185A, and Myc-HemK1. These constructs were prepared using the Gateway Cloning System (Life Technologies) and transfected into S2 cells using HilyMax (Dojindo Molecular Technologies, Inc., Japan) according to the manufacturer’s protocol.

### Fertility test

To assess female fertility, *y w* males were paired with three eclosed females of each genotype. Mating was allowed to proceed for several days in vials containing standard food. Subsequently, the flies were transferred to apple juice plates supplemented with a small quantity of moist yeast paste. The total count of eggs laid by the mated females was recorded. After a 24-hour period, the number of unhatched eggs was tallied to calculate the hatching rates. For the evaluation of male fertility in the *hemK2*-GLKD line, three *y w* virgin females were mated with *hemK2*-GLKD males over an extended duration. The fertility metrics for the males were documented in a manner analogous to that described above. Each fertility assessment was replicated across three independent biological samples.

### mRNA-seq analyses

Total RNA for each genotype was extracted in duplicate from dissected ovaries using TRIzol LS (Invitrogen) following the manufacturer’s instructions. Because *hemK2* knockdown ovaries did not contain later stages, regions post-mid oogenesis were excised from *hemK2* knockdown ovaries prior to RNA extraction. Poly(A)-tailed RNAs were enriched and purified using oligo-dT beads from the NEBNext Poly(A) mRNA Magnetic Isolation Module (New England Biolabs, USA). These samples then underwent fragmentation, reverse transcription, adapter ligation, and PCR amplification to prepare cDNA libraries using the NEBNext Ultra II Directional RNA Library Prep Kit (New England Biolabs, USA). Library preparation and next-generation sequencing were outsourced to Rhelixa Inc. (Japan), yielding approximately 13 million paired-end sequences per sample. The fastp program (Chen et al., 2018) was utilized to remove the sequences of low quality and adapter. Alignment to the *Drosophila melanogaster* reference genome (dm6) was conducted using the STAR program (Dobin et al., 2013). Read quantification at the gene level was performed with featureCounts (Liao et al., 2014). Differential expression analysis was executed via DESeq2 (Love et al., 2014), selecting upregulated (log_2_FoldChange > 1, p-adj < 0.01) and downregulated (log_2_FoldChange < −1, p-adj < 0.01) genes for further GO term enrichment analysis using PANTHER (Thomas et al., 2022).

### Monosome/Disome profiling

Ovaries were dissected in polysome lysis buffer (20 mM Tris-Cl, pH 7.5, 5 mM MgCl_2_, 100 mM NaCl) with the addition of 0.1% NP-40, 200 µg/ml emetine (Sigma, E2375), and PIC. Dissected ovaries were snap frozen in liquid nitrogen and stored at −80°C. Homogenization was performed with a plastic pestle in 500 µl of polysome lysis buffer supplemented with 0.1% CHAPS (Nacalai Tesque, Japan), emetine, and PIC. The homogenate was subjected to sequential centrifugations at 15,000 xg for 5 min, then for 15 min, both at 4°C. A small aliquot of the clear supernatant was taken for RNA quantification using the Qubit RNA HS Assay Kit (Invitrogen, Q32852). The RNA concentration was adjusted to 0.05 µg/µl with additional lysis buffer containing CHAPS, emetine, and PIC. The supernatant was treated with RNase A (1 µg/ml) at 25°C for 10 min to cleave unprotected RNA regions. After RNase treatment, 200 µl of sample was applied to a 10-50% sucrose gradient prepared in polysome lysis buffer and ultracentrifuged in a SW41Ti rotor at 30,000 xg for 3 h at 4°C. After centrifugation, gradients were fractionated from the top using a piston gradient fractionator (BioComp, Canada).

## Acknowledgments

We thank Dr Valérie Heurgué-Hamard (IBPC, Paris) and Dr. Bergmann (UMASS Medical School, Massachusetts) for generous gifts of antibody against methyl-eRF1 and UASp-P35 transgenic fly line, respectively. We acknowledge Bloomington Drosophila Stock Centre and Kyoto Stock Center for the fly stocks. Confocal images and Western blotting data were acquired with LSM900 and ChemiDoc Touch, respectively, at FBS Core Facility in Osaka University. Assistance with sucrose gradient analysis was provided by the Center for Medical Research and Education, Graduate School of Medicine, Osaka University. We appreciate the insightful discussion and suggestions from all the members of KT’s laboratory.

## Author Contributions

All authors contributed to the experiment design. XF, SR, KS, and KT performed all experiments. XF, SR and KS performed the computational analyses. XF, SR, KS, and KT wrote the paper. KS, IT, and KT supervised the project.

