## Supplementary Fig S1 v2 for "HemK2 functions for sufficient protein synthesis and RNA stability through eRF1 methylation during *Drosophila* oogenesis"

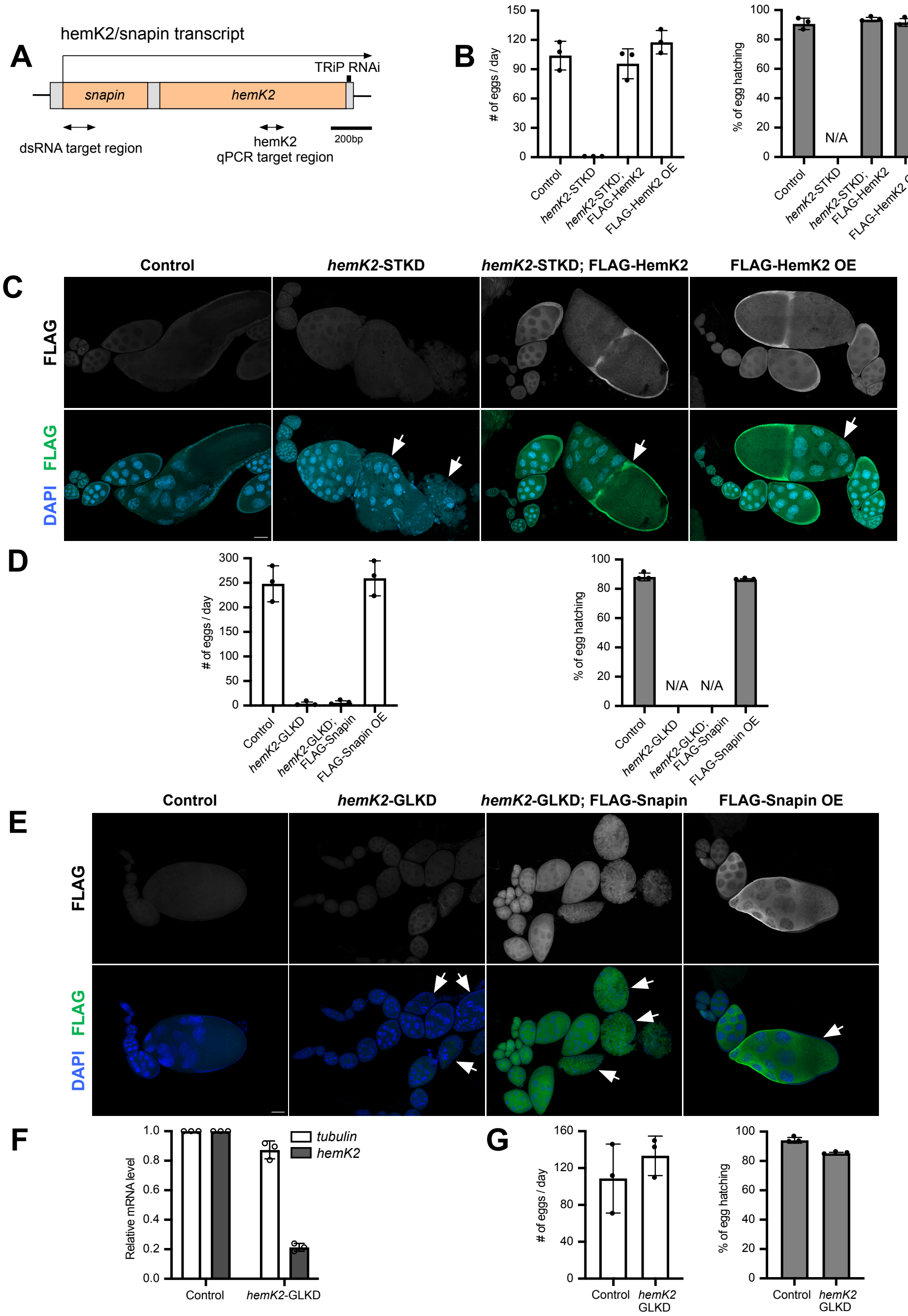


**Supplemental Figure S1. HemK2 is essential for *Drosophila* germline development.**

(A) Schematic representation of the *Drosophila hemK2/snapin* gene locus. The positions of dsRNA for *hemK2/snapin* knockdown and primers for RT-qPCR are shown below the diagram. (B) Fertility assessment through egg laying and hatching analysis. Sterility observed in *hemK2* somatic knockdown (STKD) was reversed by expression of a wild-type FLAG-tagged *hemK2* transgene under the control of the somatic driver, *traffic jam-Gal4*. The daily egg production by groups of three females is recorded (n=3 replicates). Standard deviations are shown as error bars. (C) Fluorescent immunostaining for FLAG-tagged HemK2 (green) and DNA with DAPI (blue) in ovaries corresponding to each genotype as in (B). Arrows highlight the degenerated egg chambers in *hemK2*-STKD and the progression to later stages in ovaries expressing FLAG-HemK2. Scale bars represent 50 μm. (D) Egg laying and hatching rates determined for *hemK2/snapin*-GLKD. Rescue was not achieved by expression of the *snapin* transgene alone under germline-specific drivers, *NGT40* and *nosGal4-VP16*. The quantity of eggs laid daily by three females is noted (n=3 replicates). Standard deviations are indicated as error bars. (E) Fluorescent immunostaining for FLAG-tagged Snapin (green) and DNA with DAPI (blue) in ovaries for each specified genotype as in (D). Scale bars, 50 μm. (F) qRT-PCR confirms the efficacy of the *hemK2*-GLKD in testes. The expression values are normalized to *rp49*, using *tubulin* as an internal standard. Data are presented as mean ± standard deviation from three biological replicates (n=3). (G) Analysis of egg laying and hatching. The daily egg count is from three *y w* females mated with males of the indicated genotype (n=3 replicates). Standard deviations are presented as error bars.
