## Supplementary Fig S2 v2 for "HemK2 functions for sufficient protein synthesis and RNA stability through eRF1 methylation during *Drosophila* oogenesis"

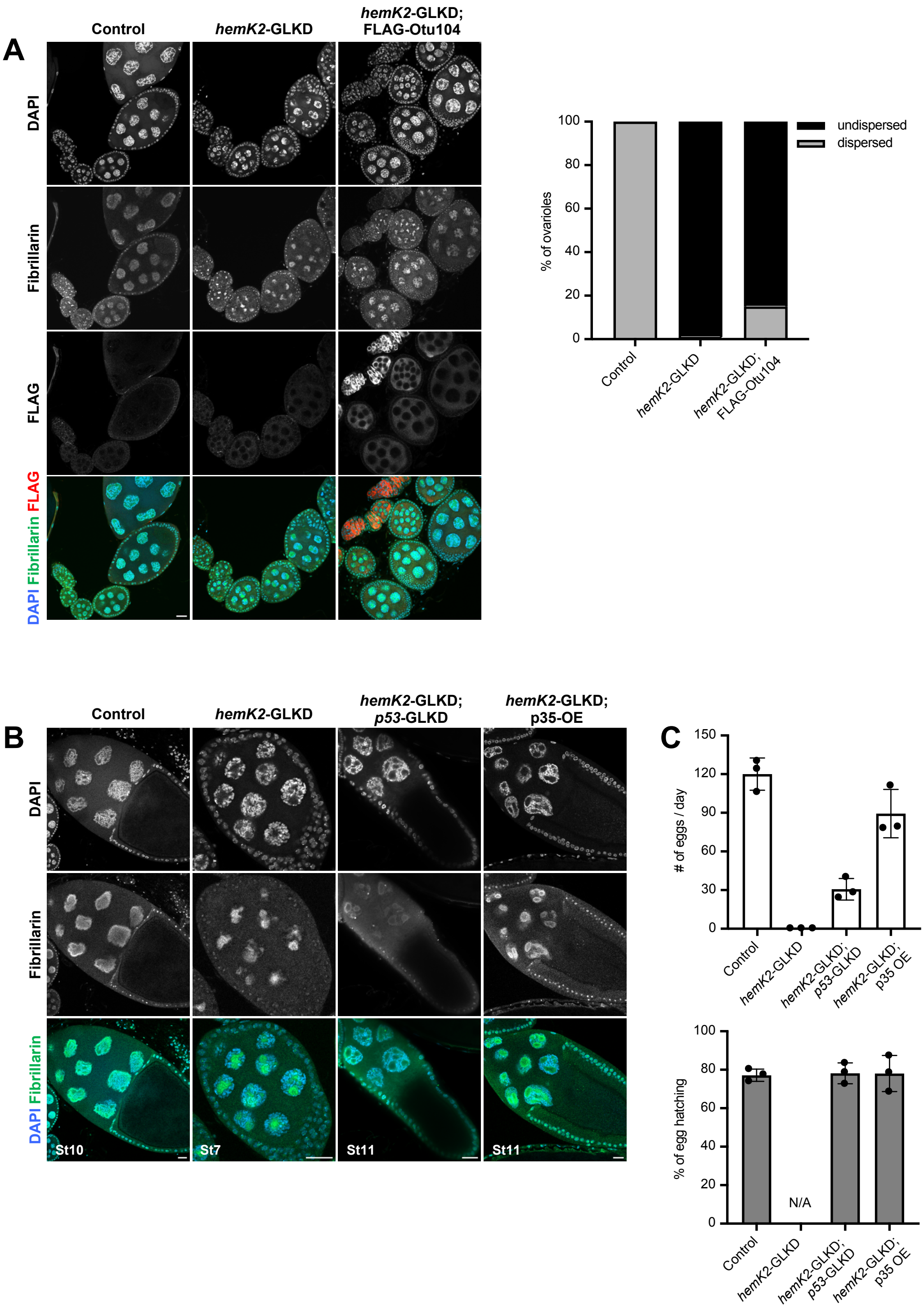


**Supplemental Figure S2. Knockdown of *hemK2* leads pleiotropic defects in germline development.**

(A) Fluorescent immunostaining for FLAG-tagged Otu104 (red), Fibrillarin (green) marking the nucleolus, and DNA with DAPI (blue) in ovaries of each specified genotype (left panel). Expression of the *otu104* transgene under the germline-specific drivers *NGT40* and *nosGal4-VP16* partially restored the chromosomal dispersion defects in *hemK2*-GLKD. Scale bars, 20 µm. The percentage of ovarioles displaying dispersed versus undispersed chromosomes was quantified for stage 6 and older egg chambers (right panel, n=100 ovarioles). (B) Immunostaining for Fibrillarin (green) and DNA with DAPI (blue) in egg chambers representing each genotype. The developmental stages of the egg chambers are determined and annotated below the images. (C) Analysis of egg laying and hatching rate for each genotype as in (B). The daily egg production was counted for groups of three females (n=3 replicates). Standard deviations are displayed as error bars.
