## Supplementary Fig S3 v2 for "HemK2 functions for sufficient protein synthesis and RNA stability through eRF1 methylation during *Drosophila* oogenesis"

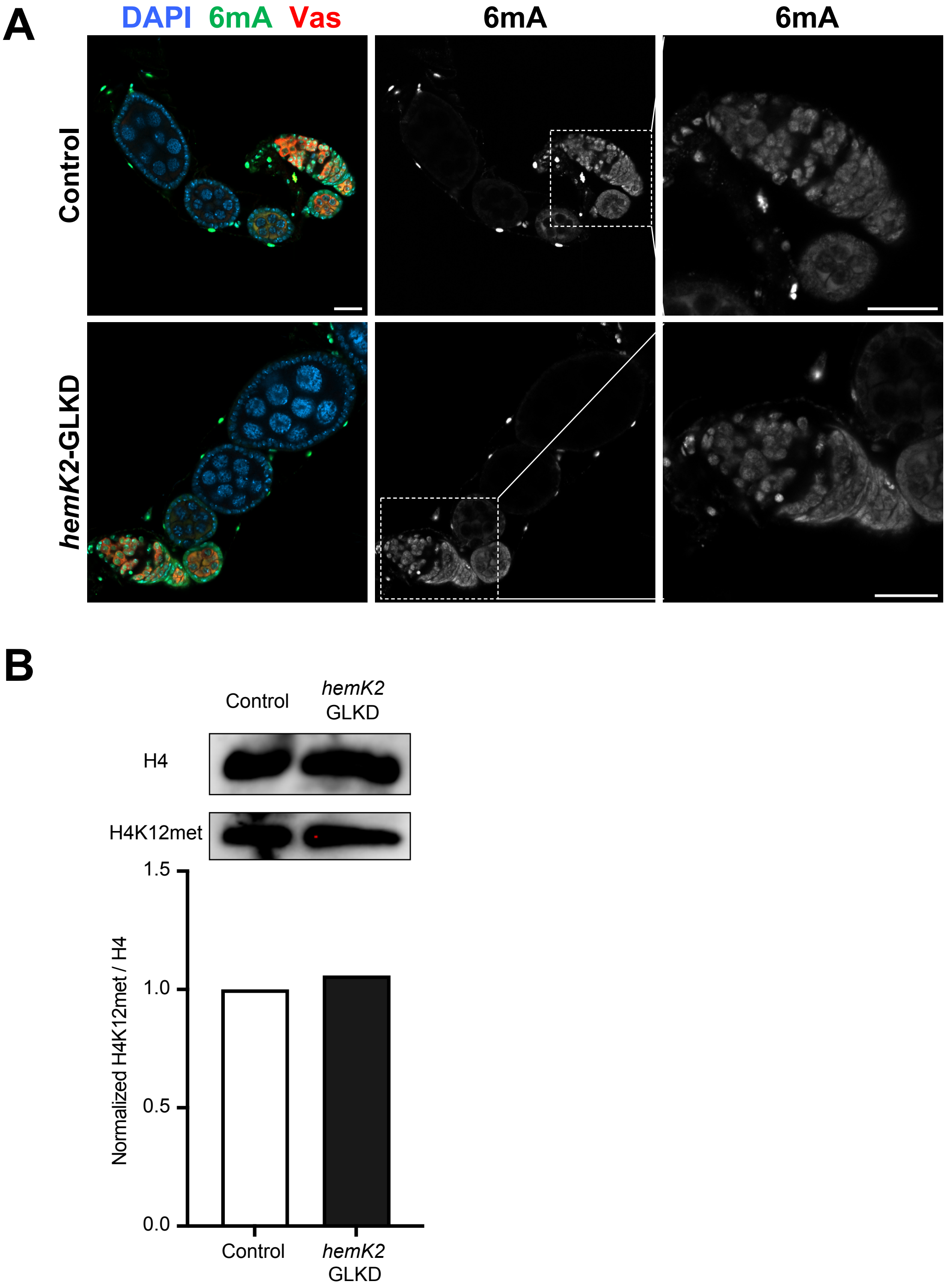


**Supplemental Figure S3. HemK2 functions for eRF1 methylation in *Drosophila*.** (A) Immunofluorescence for Vas (red) highlighting germline cells, DNA N6-adenine modification (6mA) (green), and DNA with DAPI (blue) in the control and *hemK2*-GLKD ovaries. Methylation of DNA 6mA remained consistent in *hemK2*-GLKD. Scale bars, 20 µm. (B) Western blot analysis detecting histone H4 and its methylated form at lysine 12 (H4K12met) in ovarian lysates. Quantification of the methylated histone H4K12 band intensity normalized to H4 was performed (bottom panel).
