## Supplementary Fig S4 v2 for "HemK2 functions for sufficient protein synthesis and RNA stability through eRF1 methylation during *Drosophila* oogenesis"

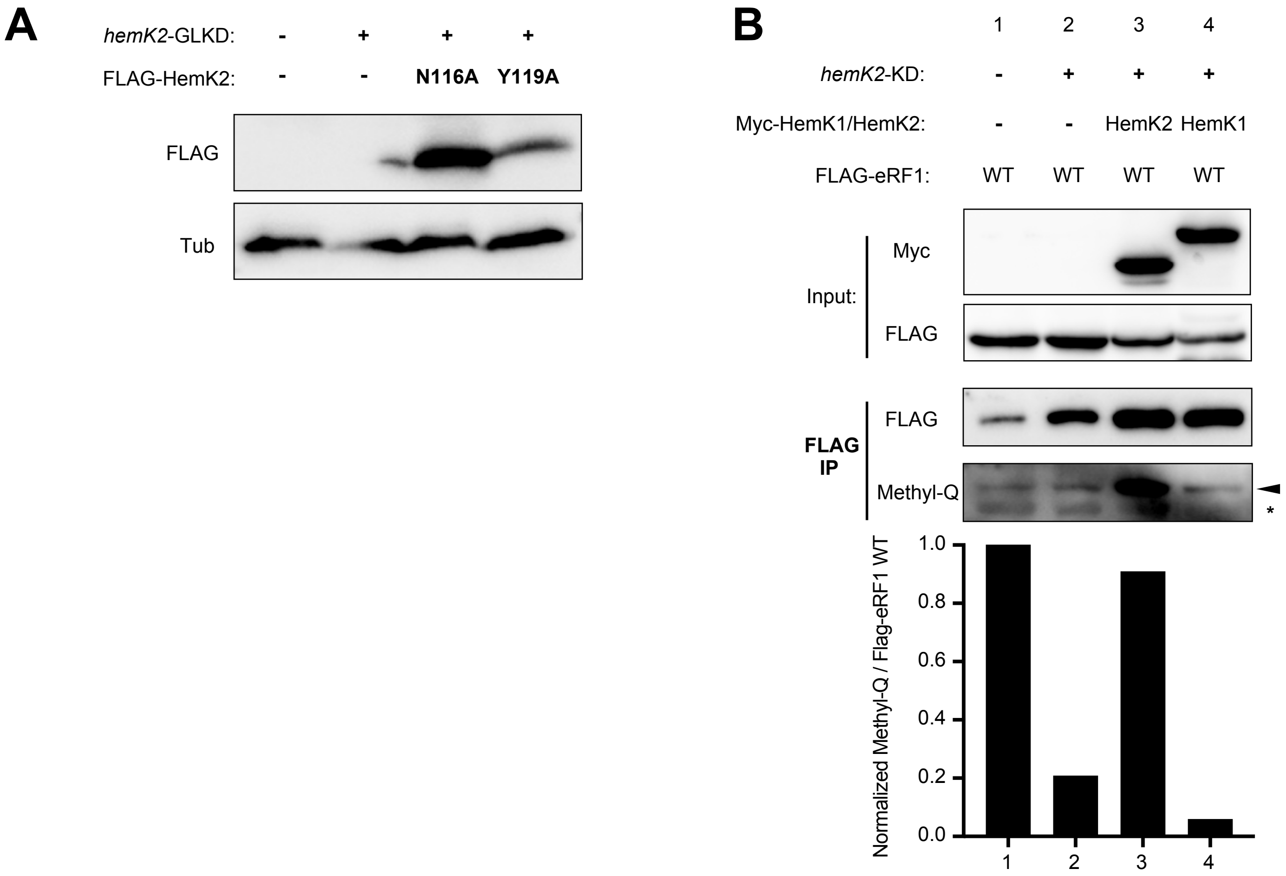


**Supplemental Figure S4. The conserved NPPY motif of HemK2 is critical for eRF1 methylation.**

(A) Western blot analysis demonstrates the expression of FLAG-tagged HemK2 N116A and Y119A mutants in ovarian lysates. Tubulin is utilized as the loading control. (B) Immunoprecipitation assay of FLAG-eRF1 conducted in S2 cells, co-transfected with Myc-tagged HemK1 or HemK2 constructs, with or without *hemK2* knockdown (KD). Western blot detection of N5-methyl-glutamine (Methyl-Q) indicates that the methylation deficit of FLAG-eRF1 due to *hemK2*-KD is restored by HemK2, but not by HemK1 expression. Intensity of methyl-Q bands, normalized to FLAG-eRF1 levels, is quantified and presented in the lower panel.
