## Supplementary Fig S5 v2 for "HemK2 functions for sufficient protein synthesis and RNA stability through eRF1 methylation during *Drosophila* oogenesis"

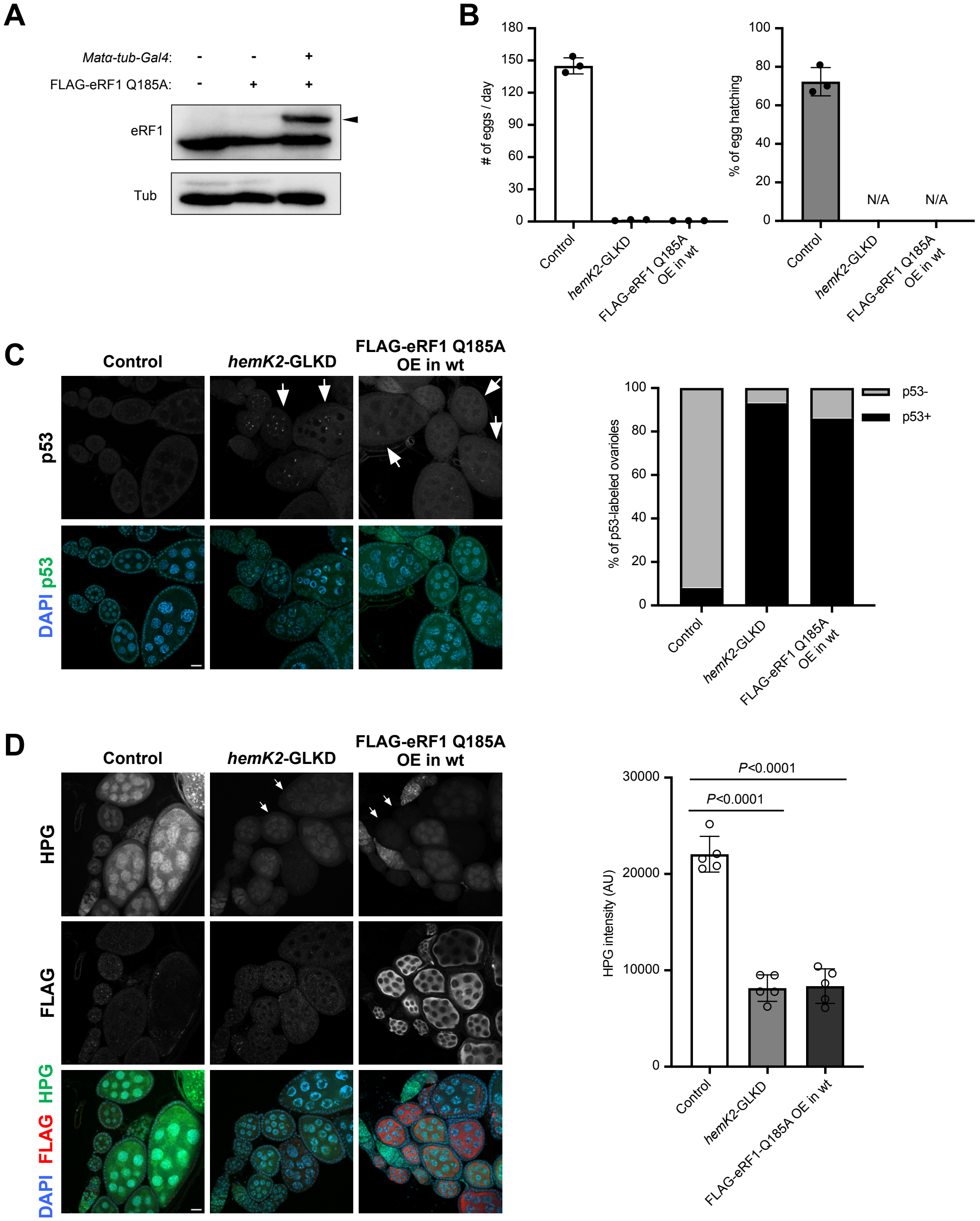


**Supplemental Figure S5. eRF1 methylation ensures proper progression of oogenesis and efficient translation.**

(A) Western blot analysis confirming the expression of endogenous eRF1 and FLAG-tagged eRF1 Q185A mutant in ovarian lysates, driven by the germline-specific *Matα-tub-Gal4*. Tubulin serves as a loading control. (B) Fertility assessment by egg laying and hatching rate for each genotype indicated. Data were collected from three females daily (n=3), and standard deviation is represented by error bars. (C) Fluorescent immunostaining for apoptosis using p53 (Green) and DAPI (Blue) for DNA in ovaries (left panel). Overexpression of FLAG-eRF1 Q185A in wildtype mimics the *hemK2*-GLKD phenotype concerning cell death. Nurse cell nuclei with concentrated p53 signals are marked by arrows. Scale bars, 20 µm. The percentage of ovarioles with or without p53 signals was quantified (right panel, n=100). (D) Homopropargylglycine (HPG) assay for protein synthesis in ovaries (left panels), with concurrent immunostaining for the FLAG epitope (Red) to visualize transgene expression. Scale bars, 20 µm. The right panel shows the quantification of HPG signal intensity across genotypes, with individual images (n=5) analyzed for statistical significance via an unpaired *t*-test. Error bars denote standard deviation.
