## Supplementary Fig S6 v2 for "HemK2 functions for sufficient protein synthesis and RNA stability through eRF1 methylation during *Drosophila* oogenesis"

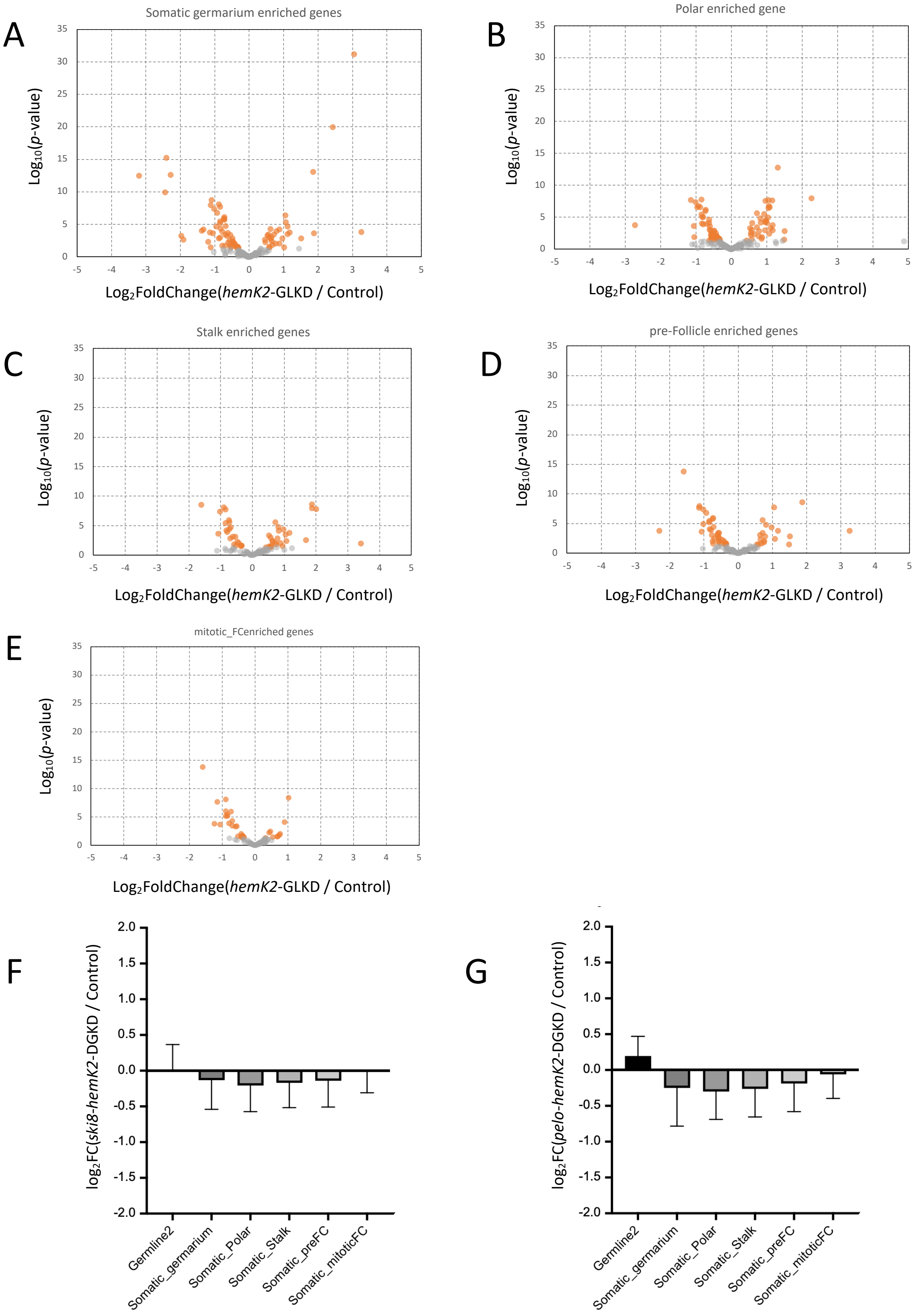


**Supplemental Figure S6. Knockdown of *hemK2* leads mRNA reduction and ribosome stalling.**

(A-E) Analysis of cell type-enriched gene expression in *hemK2*-GLKD ovaries, depicted as volcano plots. Genes significantly impacted are indicated in orange (*p*-value < 0.05, and $\log_2$FC < -1 or > 1). The cell type-enriched gene classification follows the single-cell mRNA-sequencing analysis of *Drosophila* ovarian cells (Jevitt et al., 2020). The cell types analyzed include somatic germarium (A), polar cells (B), stalk cells (C), pre-follicle cells (D), and mitotic follicle cells (E). (F, G) The $\log_2$-fold changes of cell-type enriched coding genes in *ski8-hemK2* double germline knockdown (DGKD) (F) and *pelo-hemK2* DGKD (G) ovaries in comparison to the control. Values are expressed as average $\log_2$-fold change with standard deviation.
