## Supplementary Fig S7 v2 for "HemK2 functions for sufficient protein synthesis and RNA stability through eRF1 methylation during *Drosophila* oogenesis"

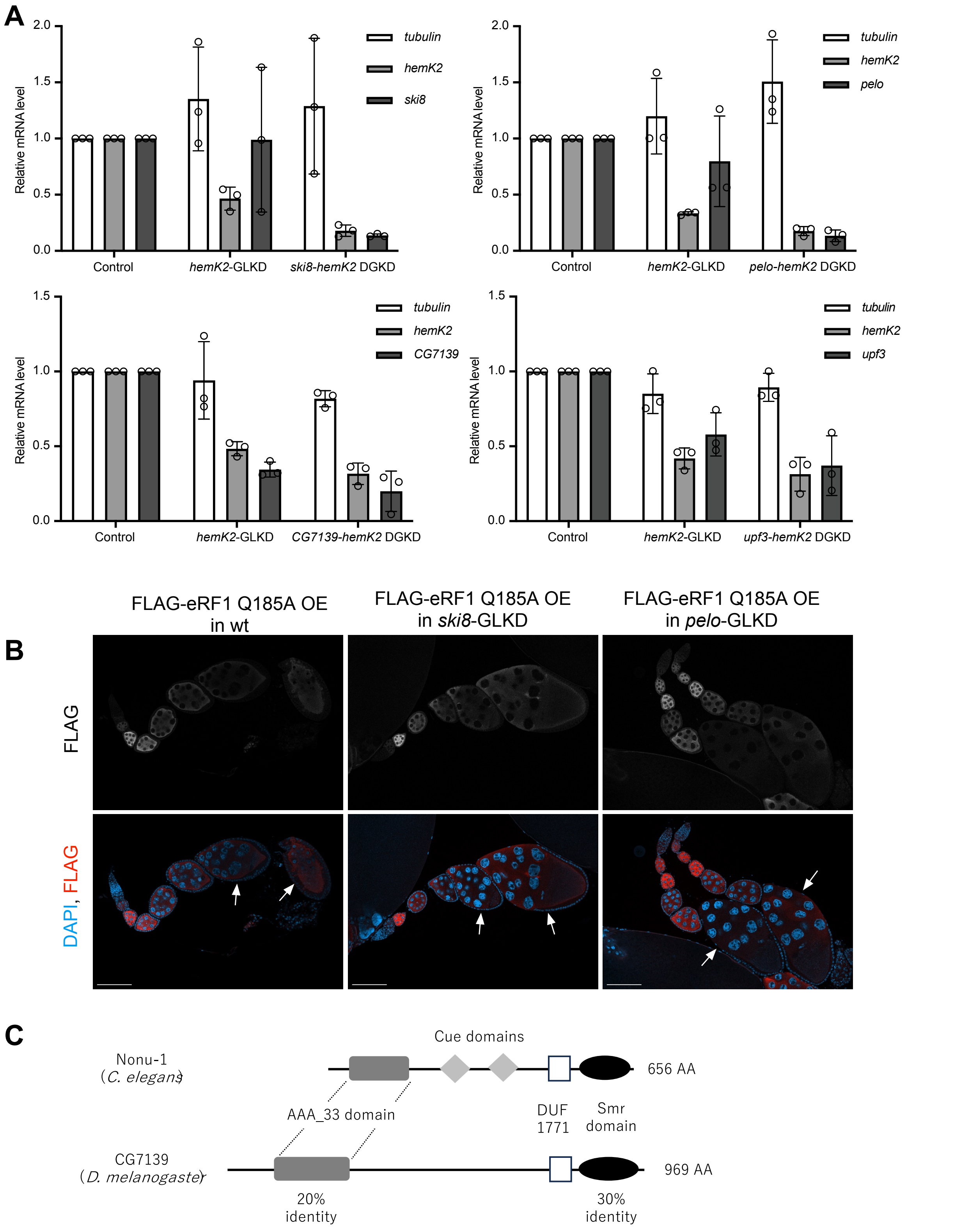


**Supplemental Figure S7. Blockage of No-Go-decay pathway restores oogenesis defects in *hemK2*-GLKD ovaries.**

(A) The knockdown efficiency of *hemK2*, *ski8*, *pelo*, *CG7139*, and *upf3* was quantitatively assessed using qRT-PCR. Standard deviation is depicted as error bars (n=3). (B) Florescent staining of ovaries for FLAG-tagged eRF1 Q185A mutant and nuclear DNA (DAPI staining, blue) in wildtype, *ski8*-GLKD, or *pelo*-GLKD genetic backgrounds. Arrows indicate degenerating egg chambers in ovaries expressing FLAG-tagged eRF1 Q185A, as well as those that were restored for progression by the concurrent knockdown of *ski8* or *pelo*. All scale bars, 100 µm. (C) CG7139 is identified as the *Drosophila* homolog of *C. elegans* Nonu-1. Sequence analysis reveals a 30% identity in the Smr nuclease domain and 20% identity in the AAA_33 kinase domain, indicating conserved functional domains.
