## Supplementary Table1 v2 for "HemK2 functions for sufficient protein synthesis and RNA stability through eRF1 methylation during *Drosophila* oogenesis"

**Supplementary Table S1. List of oligos used for generation of TRiP lines in this study.**

| Oligo label | Oligo sequence | Source |
| --- | --- | --- |
| HemK2_Fw | CTAGCAGTTAGCATCAAATTAAAGTTGTATAGTTATATTCAAGCATATACAACTTTAATTTGATGCTAGCG | Current study |
| HemK2_Rv | AATTCGCTAGCATCAAATTAAAGTTGTATATGCTTGAATATAACTATACAACTTTAATTTGATGCTAACTG | Current study |
| Pelo_Fw | CTAGCAGTAGGCAGAAACATCGAGGAGAATAGTTATATTCAAGCATATTCTCCTCGATGTTTCTGCCTGCG | Current study |
| Pelo_Rv | AATTCGCAGGCAGAAACATCGAGGAGAATATGCTTGAATATAACTATTCTCCTCGATGTTTCTGCCTACTG | Current study |
