## Supplementary Table2 v2 for "HemK2 functions for sufficient protein synthesis and RNA stability through eRF1 methylation during *Drosophila* oogenesis"

**Supplementary Table S2. List of primers used for qRT-PCR in this study.**

| **Primer label** | **Primer sequence** | **Source** |
| --- | --- | --- |
| HemK2_Fw | TGCAATGCCACTCGAAGAACT | FlyPrimerBank^a)^,  PP19952 |
| HemK2_Rv | TGACTACATAGGGCGGATTGA |  |
| Snapin_Fw | CAGCACCGTGACTTCCTTG | FlyPrimerBank^a)^,  PP11370 |
| Snapin_Rv | GAAGAGATTGGTGATGCCCTC |  |
| Poly_Fw | TCAAATGCTGGAACTCTGCTC | FlyPrimerBank^a)^,  PP21826 |
| Poly_Rv | TTATAGCCCAACCTCATGCCA |  |
| Sqd_Fw | GGTCGATGTTAAGCGTGCGA | FlyPrimerBank^a)^,  PP19425 |
| Sqd_Rv | GTCCGTCCCACTGGTTGTT |  |
| hrb27c_Fw | ACATGCCACCTAACTCTGCC | FlyPrimerBank^a)^,  PD170445 |
| hrb27c_Rv | TTGAGCACGCGAGTACATGT |  |
| hfp_Fw | ATGGAGCAGAGCATCAAGATGG | FlyPrimerBank^a)^,  PP7103 |
| hfp_Rv | GCCACACGAATTGTGTCCTC |  |
| Otu_Fw | TGCCCAAGTATGCCGGTAAG | FlyPrimerBank^a)^,  PD70094 |
| Otu_Rv | ATAGCAATTGTCCACGGGCA |  |
| Smn_Fw | CTCCGCTATTTGGGCTATGAG | FlyPrimerBank^a)^,  PP35855 |
| Smn_Rv | GGAGTCTTTGAATGACTACCAGC |  |
| Upf3_Fw | GGCGCAGTTCCTGGATCAG | FlyPrimerBank^a)^,  PP21018 |
| Upf3_Rv | ACCACCTCGCCTATGTCCT |  |
| Pelo_Fw | CCATCGCAGTGGAGAGCATAG | FlyPrimerBank^a)^,  PP18748 |
| Pelo_Rv | GCCCATCTTGACATACTGGTTCT |  |
| CG7139_Fw | GCAAACAGTGAACTCAGTAGACA | FlyPrimerBank^a)^,  PP13991 |
| CG7139_Rv: | GACAATTCGCTCTTTACGGGA |  |
| Ski8_Fw | GCACGACAGTCAGCTATGGG | FlyPrimerBank^a)^,  PP12021 |
| Ski8_Rv | GGCCTCCTTCTTATCGAAATCAA |  |
| Rp49_Fw | ATGACCATCCGCCCAGCATAC | Lin Y., et al.^b)^ |
| Rp49_Rv | CTGCATGAGCAGGACCTCCAG |  |
| Tubulin_Fw | GGTAACCGTCGAAATCAGTGTT | Lin Y., et al.^b)^ |
| Tubulin_Rv | TGGCTTTTCTGCTATACGTGTC |  |

a) Hu, Y., Sopko, R., Foos, M., Kelley, C., Flockhart, I., Ammeux, N., Wang, X., Perkins, L., Perrimon, N., and Mohr, S.E. 2013. FlyPrimerBank: an online database for Drosophila melanogaster gene expression analysis and knockdown evaluation of RNAi reagents. G3 (Bethesda), 3, 1607-1616.

b) Lin Y, Suyama R, Kawaguchi S, Iki T, Kai T. 2023. Tejas functions as a core component in nuage assembly and precursor processing in Drosophila piRNA biogenesis. J Cell Biol., 222, e202303125.
