## Supplementary Table3 v2 for "HemK2 functions for sufficient protein synthesis and RNA stability through eRF1 methylation during *Drosophila* oogenesis"

**Table S3. List of primers used for generation of dsRNA in this study.**

| Primer label | Primer sequence | Source |
| --- | --- | --- |
| T7-Snapin-Fw1 | TAATACGACTCACTATAGGGATGGATTCGGACAGCACCGTG | Current study |
| Snapin-Rv1 | CAGTGCATCCAGTTGGCCAC | Current study |
| Snapin-Fw1 | ATGGATTCGGACAGCACCGTG | Current study |
| T7-Snapin-Rv1 | TAATACGACTCACTATAGGGCAGTGCATCCAGTTGGCCAC | Current study |
